# Rich and lazy learning of task representations in brains and neural networks

**DOI:** 10.1101/2021.04.23.441128

**Authors:** Timo Flesch, Keno Juechems, Tsvetomira Dumbalska, Andrew Saxe, Christopher Summerfield

**Affiliations:** Department of Experimental Psychology, University of Oxford, Oxford, UK; CIFAR Azrieli Global Scholars program, CIFAR, Toronto, Canada

## Abstract

How do neural populations code for multiple, potentially conflicting tasks? Here, we used computational simulations involving neural networks to define “lazy” and “rich” coding solutions to this multitasking problem, which trade off learning speed for robustness. During lazy learning the input dimensionality is expanded by random projections to the network hidden layer, whereas in rich learning hidden units acquire structured representations that privilege relevant over irrelevant features. For context-dependent decision-making, one rich solution is to project task representations onto low-dimensional and orthogonal manifolds. Using behavioural testing and neuroimaging in humans, and analysis of neural signals from macaque prefrontal cortex, we report evidence for neural coding patterns in biological brains whose dimensionality and neural geometry are consistent with the rich learning regime.

## Introduction

Humans and other primates can exhibit versatile control over behaviour in rapidly changing contexts [1]. For example, we can switch nimbly between sequential tasks that require distinct responses to the same input data, as when alternately judging fruit by shape or size, and friends by gender or age [2–5]. Human studies have mapped the brain regions that exert control during task performance [6–9] or measured the processing costs incurred by task switches [10,11]. However, how the neural representations that support sequential multitask performance are acquired remains a key open question for cognitive and neural scientists [12–15].

One recently popular theory proposes that stimulus and context signals are projected into a high-dimensional neural code, permitting linear decoding of exhaustive combinations of task variables [16]. Indeed many neurons, especially in prefrontal and parietal cortex, exhibit nonlinear mixed selectivity, multiplexing information over several potentially relevant task variables [17–19], with errors heralded by a collapse in dimensionality [17]. This high-dimensional random mixed selectivity offers great behavioural flexibility because it maximises the potential for discrimination among diverse combinations of inputs, but also implies that neural codes should be relatively unstructured and task-agnostic. An alternative theory states that neural representations are mixed-selective but structured on a low-dimensional and task-specific manifold [12,13,20], with correlated patterns of firing conferring robustness on the population code [21]. Representations may adapt so that irrelevant task information is wholly or partially filtered out in ways that minimise interference between tasks [22], consistent with accounts emphasising that neural codes are sculpted by task demands [23] or through attention to scenes and objects [24]. The question of whether neural codes are task-agnostic or task-specific speaks to core problems in neural theory with widespread implications for understanding the coding properties of neurons and neural populations [25,26].

Here, we studied the dimensionality and geometry of neural codes supporting sequential multitask performance in both neural networks and the human brain. We first formalised a continuum of solutions to the multitasking problem using the framework provided by feedforward neural networks. An emergent theme in machine learning research is that neural networks can solve nonlinear problems in two distinct ways, dubbed the *lazy* and *rich* regimes, which respectively give rise to high- and low-dimensional representational patterns in the network hidden units [27–31]. In the lazy regime, which occurs when networks are initially densely wired with strong synaptic connections, the dimensionality of the input signals is expanded via random projections to the hidden layer, such that learning is mostly confined to the readout weights. In the rich regime, which occurs under small norm initializations (e.g. in initially weakly connected networks), the hidden units instead learn highly structured representations that are tailored to the task demands [27,32–34]. We used neural network simulations to characterise the nature of these solutions for a canonical multitasking setting and employ representational similarity analysis to explore their neural geometry. Subsequently, we compared these observations to BOLD data recorded when humans performed an equivalent task, and to neural signals previously recorded from macaque prefrontal cortex during context-dependent decisions [4]. In humans, we found that dorsal portions of the prefrontal cortex and posterior parietal cortex share a neural geometry and dimensionality with networks that are trained in the rich regime. This solution involves representing distinct tasks as low-dimensional and orthogonal neural manifolds, in a way that minimises interference and maximises robustness among potentially competing responses [35]. Neural signals in the two monkeys were heterogenous but we see strong support for orthogonal manifolds in one animal, with neural signals in the other strongly biased towards a single input dimension as previously reported [4,36].

## Results

We focus on a canonical paradigm involving context-dependent classification of *D*-dimensional stimuli *x*(*i, j*) ϵ *R*^*D*^ which vary along two underlying dimensions *i* and *j*, for which correct decisions depend on *i* in context *c*_*i*_ and *j* in context *c*_*j*_[2–4,14]. Healthy human participants (n = 32) categorised naturalistic (tree) stimuli, with the correct class given by *branch density* in one context and *leaf density* the other (**Fig. 1A,B, Fig S1**). These dimensions were orthogonal by design and *a priori* unknown to participants [37]. Accuracy increased with training, jumping from 64±2% to 88±2% between an initial baseline and a final test conducted in the fMRI scanner *(t*_*29*_ *= 11*.*1, p < 0*.*001*, **Fig. 1C**). Using a psychophysical model to decompose errors into distinct sources, this improved performance was due neither to a steepening of the psychometric curve (slope: *p = 0*.*120)*, nor a reduction in decision *bias* (offset: *p = 0*.*319*) although the scan session was characterised by fewer generic lapses (lapse: *Z = 3*.*5, p < 0*.*001*, **Fig. 1E**). Instead, the fitted estimation error for the category boundary fell from 27° to 7° (angular bias: *Z = -4*.*1, p < 0*.*001*, **Fig. 1E**). In a previous study [37] we quantified behavioural response patterns in this trees task by fitting a model that made choices according to the two orthogonal ground truth boundaries [37]. This *factorised* model fit better than a *linear* model that learned a single boundary for both tasks, a finding we replicate here (**Fig. 1F;** *scan: factorised > linear T*_*29*_ *= 17*.*61, p < 0*.*0001, phase x model interaction*: *T*_*29*_ *= -10*.*84, p < 0*.*0001*). In other words, despite having no prior knowledge of the tasks, or how the stimulus space was organised, participants learned over the course of training to apply the orthogonal category boundaries appropriately in each context (**Fig 1D**).

**Figure 1.**
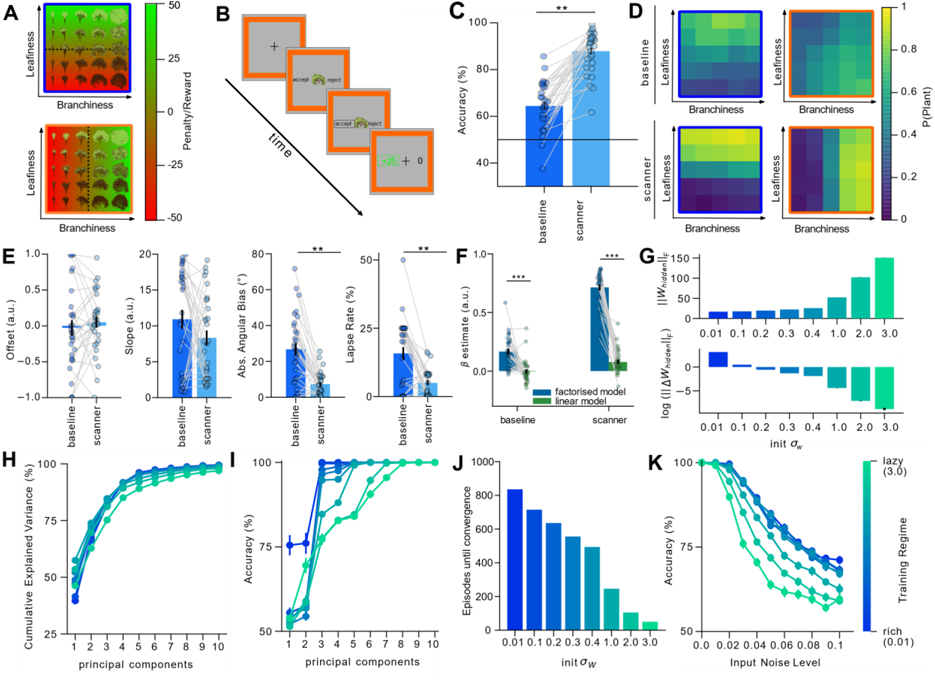
**A**. Illustration of the two-dimensional stimulus (tree) space. Each image shows the category boundary (dashed line) and reward/penalty (red-green colour) for choosing to plant in a specific context (signalled by blue frame/orange frame). **B**. Example trial sequence. Participants were asked to “accept” (plant) or reject a tree by pressing one of two buttons. The context was signalled with frame colour. Participants received rewards and penalties for planting trees. **C**. Mean accuracy improved from baseline to scan. Each dot is a participant. **D**. Choice matrices show the mean probability of choosing “plant” for each tree (defined by a level of leaf/branch density) in each context, for both the baseline (top) and scanner (bottom) sessions. **E**. Parameters of the psychophysical model between baseline and scan: offset & slope of a sigmoidal transducer, angular bias (between estimated and ground truth decision boundary), and lapse rate. See methods for details. Each dot is a participant. ** denotes p < 0.01. **F**. Fits of linear and factorised model at baseline and scan. Each dot is a participant. **G**. Norm of the weights at convergence (upper panel) and overall change in weights from input to hidden layer (lower panel) both varied with initial weight scale (x-axis and green-blue colour scale). **H**. Variance explained after the retention of 1-10 principal components of hidden layer activity (x-axis) under different initial weight scales. **I**. Network accuracy as a function of retained components. Note that the rich networks (lower initial weight scale) are more robust to compression. **J**. Episodes to convergence as a function of initial weight scale. Lazy networks converge faster. **K**. Network performance with differing levels of input noise. Rich networks are more resilient to noise.

To understand the evolution of neural codes supporting this behaviour, we trained neural networks with gradient descent to perform a simplified version of the context-based categorisation task. For simplicity, we replaced trees with stylised images (containing Gaussian blobs) that were classified according to their mean or coordinate in two interleaved contexts, signalled to the network via a unique input node. As expected from theoretical results [27,29], the norm of the weights at convergence (**Fig. 1G**, upper) and overall change in input-to-hidden layer weights over learning (**Fig. 1G**, lower) depended strongly on initial connection strengths. Networks initialised with high variance weights rapidly learned to solve the task by reading out from an approximately fixed nonlinear high-dimensional random representation (lazy regime) whereas those with low variance weights converged more slowly, but exhibited strong representation learning in the input-to-hidden weights (rich regime). Thus, the final representations were lower dimensional under rich learning, with just 6 (9) principal components needed to explain 95% of the variance under rich (lazy) learning (**Fig. 1H)**. Critically, however, the rich regime proved more tolerant to a challenge that reduced the dimensionality of hidden unit activity: only 3/6 components were needed to maintain ceiling performance, whereas 8/9 were required under lazy learning (**Fig. 1I**). Although learning was up to 10 times faster in the lazy regime (**Fig. 1J**), the highly structured representations acquired during rich learning conferred robustness, also making performance more tolerant to the addition of Gaussian input noise (**Fig. 1K**). In other words, these solutions offer complementary costs and benefits for representation learning (speed vs. robustness) of task-related variables.

Next, we used representational similarity analysis (RSA) and multidimensional scaling (MDS) to visualise the neural geometry of the network hidden units at convergence under either regime (**Fig. 2A**). Focussing on the minimum and maximum norm solutions, during lazy learning the similarity is mostly driven by the structure of the input space (including the task context) (upper panel); this is expected because the input weights remain close to their initial values and random high-dimensional projections approximately preserve distances between inputs [38]. However, during rich learning hidden unit activity varies with context: in *c*_*i*_, neurons code for dimension *i* but not *j*, with the converse true for *c*_*j*_. In other words, task-irrelevant features were filtered out in each context, transforming the neural “grid” into two manifolds, each coding for a task-relevant dimension. Specifically, each context has a compressed and uncompressed axis, forming a rectangle in the plane, and we hereafter call the geometry “orthogonal” when the respective compressed and uncompressed axes are perpendicular across tasks. Thus, the network learned to project the data on a low-dimensional embedding space, in a way that might minimise intrusions from irrelevant features in each context (lower panel)[35]. This was confirmed by fitting model representational dissimilarity matrices (RDMs) that encode a grid or orthogonal pattern to the hidden units at convergence: the grid model fit best for high-norm (lazy) solutions and the orthogonal model fit best for low-norm (rich) solutions **(Fig. 2D**).

**Figure 2.**
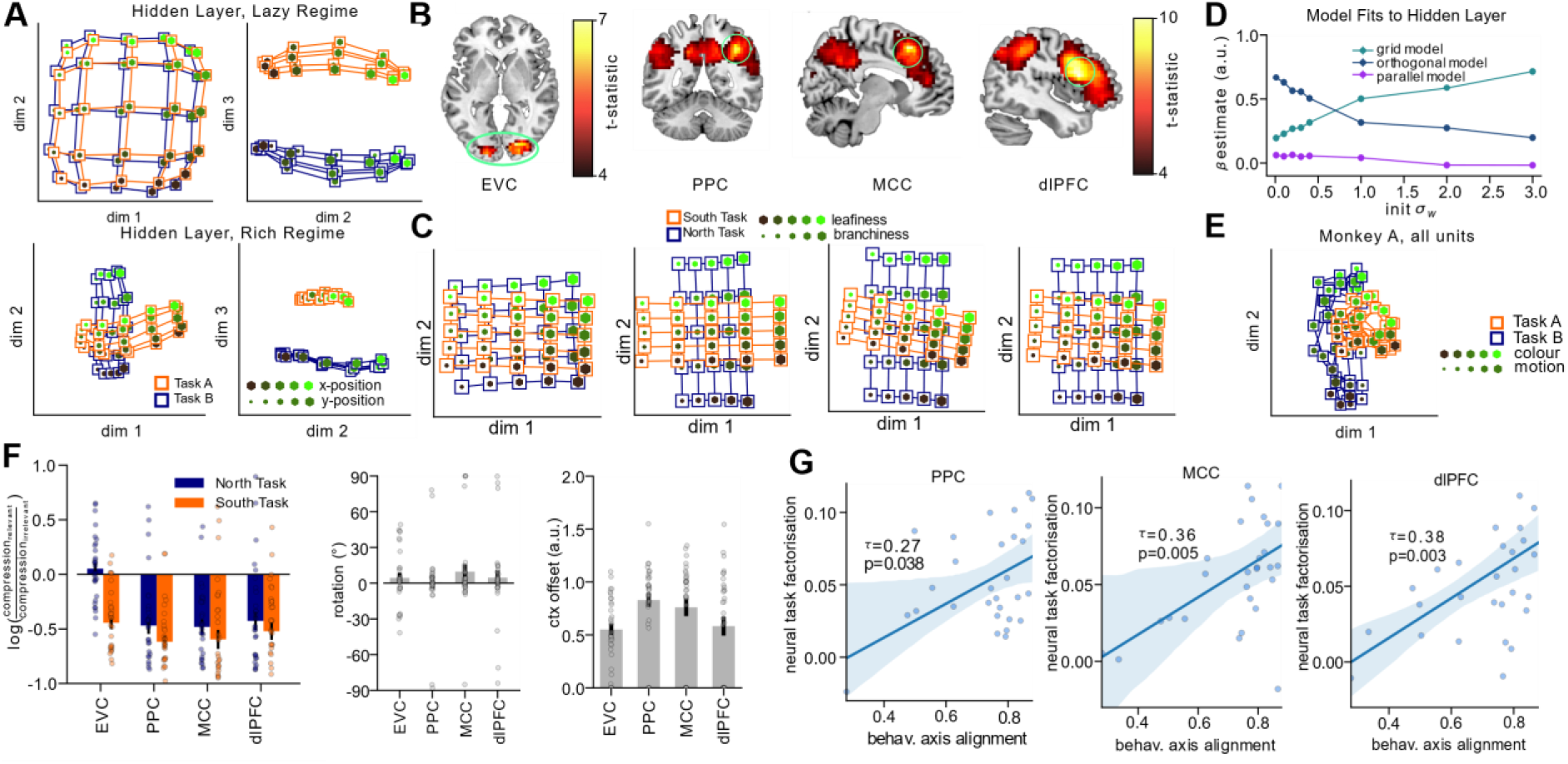
**A**. Three-dimensional representation of hidden layer representations for each stimulus feature (x- and y-position, dot colour and size) in each context (connecting lines, orange and blue). Top panels: lazy regime, bottom panels: rich regime. In the rich regime, note how the compression along irrelevant dimensions leads to the emergence of “orthogonal” manifolds, in which task-relevant stimulus information is encoded along orthogonal axes. **B**. Left panel: voxel regions where neural similarity patterns matched the grid RDM. Right panel: voxels where neural similarity patterns matched the orthogonal RDM. All data are corrected for multiple comparisons. **C**. Low-dimensional projections of fMRI data from within ROIs taken from visual, parietal and frontal regions, reconstructed from coefficients of regression model **D**. Fits of RDMs encoding grid, orthogonal and parallel representational schemes to the neural network data as a function of initial weight scale. The orthogonal model (dark blue line) fits best in the rich regime, and the grid model (cyan line) fits best in the lazy regime. **E**. Same as A and C but for data from *monkey A*. Stimulus features are now colour and motion; data from Mante et al 2013 [4]. **F**. Data from parametric RDM fits. Compression, rotation and context offset in each region from best-fitting RDM characterised by parametrically varying expansion/contraction of representation on relevant/irrelevant dimension (left panel), context-dependent rotation of the stimulus axes from native space into the reference frame of the response (i.e. from orthogonal to parallel model, mid panel) and separation between contexts (right panel). **G**. Correlation between neural task factorisation (fits of orthogonal model to neural data) and behavioural axis alignment (fits of factorised model to choice matrices). Each dot is a participant.

How, then, are task representations structured in biological brains? Our simulations furnished predictions about the neural geometry we should expect to see in BOLD data acquired during the final phase of our experiment. Univariate tests replicated standard findings including the heightened BOLD signal in PFC on context switch relative to stay trials (**Fig. S2A,B**), and the correlation between BOLD signal and decision certainty in posterior parietal [39] and medial orbitofrontal cortex [40] **(Fig. S2C,B)**. However, to investigate neural geometry, we once again turned to a more powerful multivariate analysis of the activity patterns (RSA). We used model RDMs encoding grid, orthogonal and control patterns to predict brain activity. Crucially, we observed strong correlations with the *orthogonal* model in three major foci: the dorsolateral prefrontal cortex (dlPFC; *t*_*30*_*=9*.*79, p<0*.*001 corrected, peak [46 14 24]*), the mid-cingulate cortex (MCC; *t*_*30*_*=9*.*51, p<0*.*001 corrected, peak [8 21 49]*) and the posterior parietal cortex (PPC; *t*_*30*_*=8*.*87, p<0*.*001 corrected, peak [39 -45 45])* **Fig. 2B)**. A similar effect was observed in a left prefrontal region for which the univariate analysis had revealed that it was sensitive to context switches, but the fit of the orthogonal model did not differ between switch and stay trials (**Fig. S3**). In early visual regions, neural data RDMs were best predicted by a model in which dissimilarities depended mainly on branch density (*t*_*30*_*=6*.*98, p<0*.*001 corrected, peak [22 -84 -3]*) but no other models explained a significant amount of variance in the neural RDMs. Thus, neural codes were largely structured as predicted by rich learning, with representations in each context projected onto orthogonal neural axes that are elongated along the relevant feature dimension and compressed along the irrelevant feature dimension.

We also used RSA in conjunction with a parametric model-fitting approach conducted on independently defined ROIs for dlPFC, MCC and PPC. Rather than fitting models encoding extremes of compression, rotation, and context separation, now we built RDMs by varying these factors continuously, visualising the parameters that best fit the neural data in each region. This confirmed that the neural code was compressed along irrelevant but not relevant dimensions and remained in the naïve (input) space rather than being rotated into the frame of reference of the response (**Fig. 2F**). When we used MDS to visualise the best-fitting model RDMs for each region, the task-specific encoding of relevant dimensions along orthogonal manifolds in dorsal stream regions of interest can be clearly seen **(Fig. 2C)**. Finally, in neural networks rich learning is characterised by a low-dimensional neural code; by systematically removing components from the data using PCA on the BOLD patterns within each candidate ROI, we were able to show that reliable correlation with the orthogonal manifolds RDM required just two components in each region of interest and that there was no measurable benefit in maintaining more than 6 PCs in total (**Fig. S4**). In other words, the neural representations span a low-dimensional subspace focused on task-relevant stimuli, as predicted by rich learning.

Next, we attempted to link these neural patterns to behaviour. The factorised model that was fit to human choices to quantify the extent to which these were aligned with the ground truth category boundaries yields an “axis alignment” score for each participant, which was correlated with the orthogonality of neural task representations across the cohort in PPC (*Kendall’s tau a = 0*.*27, p=0*.*038*), MCC (*Kendall’s tau a=0*.*36, p=0*.*005*) and dlPFC (*Kendall’s tau a = 0*.*38, p=0*.*003*; **Fig. 2G**). In other words, the decisions of participants with more factorised neural representations respected more orthogonal category boundaries.

BOLD data offers at best an indirect window on neural coding, so we additionally capitalised on a freely available dataset describing single neuron activity in frontal eye fields (FEF) whilst macaques performed an equivalent context-dependent decision task on stimuli with varying colour and motion coherence [4,36]. We focus on the results from monkey A, because our analyses (and those reported previously) indicate that FEF neurons recorded from monkey F were only very weakly sensitive to motion even when it was decision-relevant [36]. First, we built a pseudopopulation from all the recorded neurons and plotted its neural geometry in 2 dimensions. This revealed two orthogonal manifolds, each coding for one of the two task-relevant axes, just like in the BOLD data and predicted by neural networks trained in the rich regime (**Fig. 2E**). Indeed, when we fit the candidate RDMs used above to this dataset, the orthogonal RDM fit best for monkey A; an RDM coding for colour alone fit best for monkey F (**Fig. S4**). We also tested dimensionality of these neural geometries using a similar approach as above; the ability to decode orthogonal manifolds dropped sharply when fewer than 3 components were retained, suggesting that directions of highest variance were aligned with the task-relevant dimensions of context, colour and motion (**Fig. S5**). This analysis suggests that the orthogonal manifolds identified with the RSA lie embedded in a very low-dimensional manifold and indicates that the effect observed in human BOLD generalises across species and recording methods.

How does this neural coding scheme prevent interference among tasks? In the neural network model, we reasoned that orthogonal manifolds could emerge if the weights linking each context unit to the hidden layer were anticorrelated. Anticorrelated weights ensure that distinct subsets of hidden units are active in each context, as neurons which receive negative net input in one context (and which therefore are inactive due to the rectified linear (ReLU) activation function) will receive positive net input (and be active) in the other. By wiring only the task-relevant stimulus dimension to the active population in each context, information along the irrelevant dimension is thus effectively zeroed out by the nonlinearity, creating an independent subspace for each task (**Fig. 3A**). This would allow the network to factorise the problem, encoding the task-relevant information in a way that avoids mutual interference (**Fig. 3B-D**).

**Figure 3.**
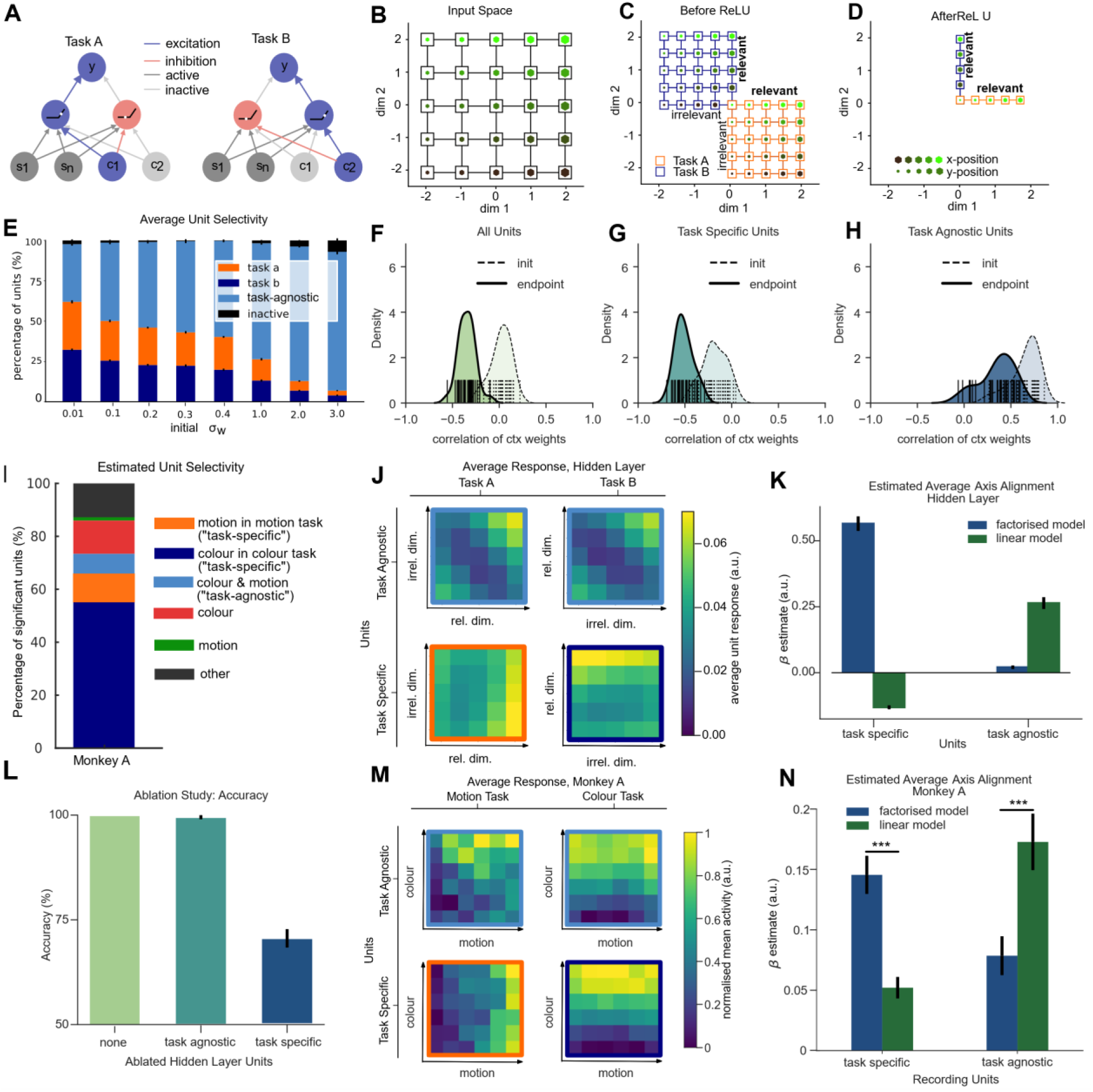
**A**. Schematic illustration of how opposing weights from two context units leads to learning two unique subspaces. Red and blue arrows show positive and negative weights from context units, which control the sign of the net inputs in the hidden layer, so that stimuli are effectively processed by different hidden units in each context. **B-D**. Schematic illustration but in neural state space. B. Shows similarity structure among input stimuli with no context modulation. C. Shows the similarity structure in the hidden layer net input (before ReLU). Note the separation between contexts. D. After the ReLU, “inhibited” (below-zero) inputs are removed, leaving two orthogonal manifolds. **E**. Task and stimulus selectivity in the neural network as a function of initial weight scale. **F**. Distribution of empirically observed correlation coefficients among context unit weight vectors in the neural network. **G-H**. Same as F but separated out by “task-selective” and “stimulus-selective” units as defined in E. Note the anticorrelation in task-selective units (and overall). **I**. Distribution of selectivity of single neurons in monkey A, using the same criteria as in E. **J**. Hidden unit selectivity for each relevant and irrelevant stimulus feature in each context. Note that task-selective units (lower panels) are mostly sensitive to relevant vs. irrelevant dimension whereas stimulus-selective units code for an interaction between features. **K**. Quantification of results in J using fits of linear vs. factorised model. The factorised model fits best to task-selective units, and the linear model to stimulus-selective units. **L**. Results of ablation study. Ablating task-selective, but not stimulus-selective units is detrimental to performance. **M**. Same as J, but for example neurons from monkey A. **N**. Same as K, but for monkey A.

This theory makes a number of testable predictions. Firstly, it implies that most neurons should be mixed selective, responding to combinations of stimuli and task variables. Indeed, we observed that a large proportion of neurons were mixed selective regardless of regime. Secondly, however, it implies that this mixed selectivity should be structured in the rich regime, with most units in the hidden layer responding specifically to the combination of task-relevant stimulus dimension and task. Indeed, we observed that up to ∼60% of hidden units responded exclusively under one task or the other during rich learning (**Fig. 3E**). Interestingly, when we conducted a comparable analysis for non-human primate (NHP) data, we found that the majority (∼65%) of significantly responsive units were also selective to either colour in the colour task or motion in the motion task, although there was strong bias towards the colour task (**Fig. 3I**). Thirdly, the theory predicts that in neural networks the context weights should be anticorrelated. This is indeed the case on average in the rich regime (**Fig. 3F**) and especially for the majority of task-selective neurons (**Fig. 3G**), which became anticorrelated as training progressed. In contrast, those neurons that converged to being task-agnostic were those that received strong, positively correlated input from two context units at random initialisation, and this input remained positively correlated after training (**Fig. 3H**). It thus seems likely that the initial sign of the connections from the context units to each hidden unit determines whether it is destined to be a task-agnostic or task-specific unit during training. We cannot test this in NHP data, but we can compare the response profiles of neurons defined as task-agnostic and task-specific in both model systems, revealing how their responses vary with stimulus input in either context. The theory predicts that task-specific units show a coding preference for relevant feature dimensions (with irrelevant features mapped onto units which are deactivated by the ReLU). This is exactly what is seen in both the neural network (**Fig. 3J,K** *factorised model > linear model: z = 4*.*781, p < 0*.*0001, d = 0*.*873*) and the NHP data, where the responses of task-specific units are aligned to the two choice axes (**Fig. 3M,N** *factorised model > linear model*: *z = 4*.*916, p < 0*.*0001, d = 0*.*739*). By contrast, in neural networks the remaining ∼35% of active units coded for a residual policy which collapses across both contexts (“task agnostic”), resembling the *linear* model described above (**Fig. 3J,K** *linear model > factorised model z = 4*.*781, p < 0*.*0001, d = 0*.*873*). The same task-agnostic response patterns were observed in NHP neurons that responded significantly to stimuli but did not differentiate substantially between dimensions **(Fig. 3M)**. Just as in the neural network simulations, responses of these single units were best explained by the linear model (**Fig. 3N** *linear model > factorised model: z = 4*.*076, p < 0*.*0001, d = 0*.*732*). A final prediction of this theory is that in the rich regime, performance depends critically on the task-specific (and axis-aligned) neurons but not on those displaying task-agnostic selectivity. In the neural network, we thus conducted an ablation study in which the output of either the task-agnostic or task-specific neurons was set to zero at evaluation. Performance was unimpaired by the loss of task-agnostic units but dropped to ∼70% after task-specific units were removed, consistent with the use of a single linear boundary across the two contexts (**Fig. 3L**). Together, these findings support a model of context-dependent decision-making whereby the network learns to gate information into orthogonal subspaces in the hidden units (of a neural network) or prefrontal cortex (of humans and NHPs), in a way that minimised mutual interference. This scheme emerges when context input signals are anticorrelated.

## Discussion

The work described here makes three distinct contributions. The first is to formalise solutions to the learning of a canonical context-dependent classification paradigm using a feedforward connectionist (or “deep learning”) framework [41,42]. We do this by drawing upon recent work in machine learning research, which distinguishes among the learning regimes which occur when deep networks are initialised with strong, dense connections (high norm weights; lazy regime) compared to weak connections (low norm weights; rich regime) [16,17,27–32]. We derive predictions from these regimes for the context-dependent classification task, a paradigm that has been well-studied before using both single neuron electrophysiology [4,36] and neuroimaging [2] methods.

The second contribution is to assess these predictions using behavioural testing and functional neuroimaging in human participants, and reanalysis of a dataset recorded from macaque monkeys performing an equivalent task. In humans, we find that over the course of training, participants learned about the structure of the stimulus space and correctly inferred the orientation of the two category boundaries. After training, we observe a stylised neural geometry in the parietal and prefrontal cortices that closely matches the predictions of the “rich” regime, whereby stimuli are projected onto orthogonal subspaces on a low-dimensional manifold. A similar pattern was observed in the NHP data. Together, these data speak to a debate about whether humans and other primates learn to solve complex tasks by forming high-dimensional (and task-agnostic) or low-dimensional (and task-specific) neural codes and offer striking evidence for comparable coding principles in humans, non-human primates and artificial neural networks.

The third contribution is an insight into the computational principles that allow the context-dependent decision task to be solved. We show that a combination of anticorrelated context inputs and ReLU (or ReLU-like) nonlinearities allows the network to effectively learn to gate task information according to context. This allows us to predict how mixed-selective neurons code for relevant and irrelevant features in both neural networks and NHPs, and to anticipate the effects of silencing task-agnostic vs. task-specific neurons on performance. We note that for the NHP task, where inputs arrive over time, our simple theory models the representation at late times after stimulus presentation. Adding recurrent connectivity yields a model exhibiting a “late selection” mechanism and fixed stimulus input directions across contexts, two key hallmarks identified in prior analyses [4,36](see Fig. S8 and Supplementary Methods).

There has been a recent resurgence of interest in neural networks (or “deep learning models”) as computational theories of biological brains [41,42]. A common approach is to use linear methods to examine similarities between the representations formed in biological systems (e.g. multi-neuronal or multivoxel patterns) and in the hidden units of deep networks. One corollary of our findings is that the relationship between representations formed in biological and artificial networks can critically depend on the variance of the weights at initialisation. For example, when the initial weight scale is large, the similarity structure of encoded representations will closely match their input structure. This is what we saw in BOLD data from visual cortex (in our case, a more “grid-like” pattern, with higher sensitivity to variations in shape than in colour). This may partly explain why previously reported improvements in model fit from trained to untrained networks tend to be relatively modest, as if the visual cortex mainly recapitulates the input data through random high-dimensional projections [43,44].

In our data, the nature of the neural code observed in parietal and prefrontal cortex, however, was very different. Here, task-irrelevant features were compressed in each context, converting the neural “grid” into orthogonal manifolds, each coding for a task-relevant axis. This is quite striking, because conflicting reports have suggested that task-irrelevant information is retained or discarded during context-dependent decision-making [2,4]. More generally, the diverse representational structure that can emerge in the rich and lazy regimes, and its variable mapping to the brain, may shed light on why emerging representation structure can be heterogenous in trained neural networks [45].

Previous analyses of single cell data from macaque prefrontal cortex have emphasised that neural selectivity is mixed, and representations are high dimensional, in seeming contradiction to the findings reported here [16,17]. One possibility is that over prolonged training, the dimensionality of neural representations is tailored to the transfer demands of the paradigm [46]. Structured, low-dimensional representations may be favoured in settings where information can be shared across tasks or stimuli, such as our trees task, where all stimuli were unique, but sampled from the same underlying generative process, hence permitting generalisation of latent features across tasks. By contrast, high-dimensional neural codes may emerge by preference in tasks with minimal need for generalisation, such as recall and recognition of a small set of unrelated images [17,47]. Indeed, our “rich” neural networks were more tolerant to degradation through compression and/or input noise than those in the lazy regime. However, the relationship between the generalisation ability of the two regimes described here remains an open question.

At first glance, our findings might appear to diverge from previous analyses of the same data, in that we emphasise that irrelevant information is at least partly compressed in FEF [4,36]. However, our analysis of the NHP data focussed on a relatively late epoch (300-600 ms post-stimulus). In fact, when we repeated the model-based RSA separately for early, middle and late time windows following stimulus onset, we found that representations were more grid-like early on (encoding of both feature dimensions) but became highly task-specific in the second half of the trial (**Fig. S8a**). Crucially, we can explain this temporal evolution of task representations with an extension of our gating theory that incorporates recurrence into the neural network model (**Fig S8b**). Under this account, feature-selective units keep integrating motion/colour information throughout the stimulus presentation period, but the irrelevant dimension is integrated at a slower rate, giving rise to a gradual progression from grid-like to orthogonal representations. In the following delay-period, the context cue continues to act as inhibitory bias on the unit encoding irrelevant features, gradually supressing its activity just enough so that by the time of a response, only task-relevant information is preserved, leaving a fully orthogonal and task-specific representation (**Fig S8c**). When we visualised the geometries separately for early, middle and late windows within the stimulus interval, we observed a similar temporal evolution from grid-like to more orthogonal representations in both the RNN and monkey recordings (**Fig S8d**).

Another recent paper has emphasised that the neural geometry for distinct tasks in macaque PFC can become aligned along parallel manifolds, with representations for common action/ outcome associations aligned in neural space [48]. An equivalent effect in our paradigm would be that tree representations are rotated into a frame of reference of “plantworthiness” – whether the tree should be accepted for planting or not – which we tested with a “parallel model” RDM but failed to find evidence for in either neural data or the network hidden units. One important difference in our work is that in order to separate decision and motor activity, in the fMRI study we varied the motor contingencies from trial to trial, meaning that there is no real benefit to representing the decision directly in the response frame in our task. In fact, further neural network simulations revealed that in a two-layer neural network, orthogonal representations dominated in the first hidden layer, but more parallel representations emerged in the subsequent layer, more consistent with the findings of [48] (**Fig. S7**). We take this to imply that in a task where response contingencies were not randomised from trial to trial, we might see parallel representations emerge in a putative downstream stage – for example premotor cortex – but this contention remains to be tested.

Taken together, our findings suggest striking similarities between multi-task learning in biological and artificial neural networks and indicate that the human brain has evolved a coding scheme that minimises representational overlap between consecutively learned tasks, similar to the one adopted by a neural network trained in the rich regime on interleaved data.

## Acknowledgements

This work was supported by generous funding from the European Research Council (ERC Consolidator award to C.S. and Special Grant Agreement 3 of the Human Brain Project) a Sir Henry Dale Fellowship to A.S. from the Wellcome Trust and Royal Society (grant number 216386/Z/19/Z). and a Medical Science Graduate School Studentship to T.F. (Medical Research Council and Department of Experimental Psychology). A.S. is a CIFAR Azrieli Global Scholar in the Learning in Machines & Brains programme.

## Methods

### Human Behavioural / fMRI Experiment

#### Participants

A total of 32 participants (mean age 24.44y, 31 right-handed, 21 female) with no history of neurological or psychiatric disorders were recruited from a participant pool at the University of Granada. One participant was excluded from the analysis due to equipment failure during the scanning session, leaving 31 participants for the fMRI analysis. For another participant training data was not recorded due to disruption of their internet connection, leaving 30 participants for all behavioural analyses. All participants gave written informed consent prior to taking part in the study. The experiment received approval from the ethics board of the University of Granada. Participants were compensated for their time with 38€. The experiment consisted of several sessions completed on three successive days (**Fig S1a**). All participants completed a pre-screening study on day 1 that assessed their eligibility for the main experiment. The main experiment consisted of a browser-based training session on day 2, and a refresher and scanning session on day 3, which took place at the fMRI institute of the University of Granada.

#### Stimuli

Participants performed a virtual gardening task for which they had to discover rules that determined growth success of tree stimuli in two different gardens. Trees were generated by in house-code [link] and were built to vary parametrically in five discrete steps along two different dimensions, the density of leaves (“leafiness”) and the density of branches (“branchiness”), yielding 25 unique class. We generated multiple stimuli per level of leafiness and branchiness and sampled these exemplars randomly without replacement for training and test sessions at the level of individual participants so that no physical stimulus was presented twice during the experiment.

#### Pre-Screening Session (Day1)

We previously showed that learning is mediated by an *a priori* tendency to factorise tree space into dimensions of leafiness and branchiness [37]. To measure this prior in our participants we first used an online task in which participants moved tree exemplars within a circular open arena via drag and drop on the screen, attempting to arrange them so that distance between trees was proportional to their perceived dissimilarity (**Fig. S1b**). Participants completed six arrangement trials of 25 trees, with trees sampled from the whole 5×5 grid of branchiness and leafiness on each trial. At the beginning of each trial, the trees were randomly arranged in an attempt to minimise other sources of bias. The allocation of exemplars to trials was randomised across subjects. We correlated the dissimilarity matrices derived from the arrangements with a model matrix that represented a perfect grid-like arrangement to compute a “grid score” for each participant. We planned to exclude participants who failed to reach the median grid score reported in the previous study where participants were recruited online [37], but no participants met this criterion (**Fig. S1c**).

#### Training Session (Day2)

On day 2, participants took part in an online training session in which they learned to perform the task. On each trial participants first viewed a cue indicating the context (or “garden”), which was a blue or orange rectangular frame presented for 1000ms. Next, a tree was displayed for 1500ms within the frame, together with the response contingencies (“plant” or “don’t plant”) which were indicated by left and right arrow buttons on either side of the tree stimulus. These contingencies (i.e. whether “plant” was mapped onto the left or right button) were varied randomly from trial to trial. The stimulus and response interval was always set to 1500ms. If a response provided within this interval was highlighted by a rectangle drawn around the chosen option (“plant” vs “don’t plant”). Participants were asked to learn to plant trees that grew successfully. Tree growth success depended on leafiness in one context and branchiness in another and was signalled by a numerical reward, ranging in five steps from -50 to +50. For example, for a given participant, trees occurring within the orange frame might grow successfully if they had fewer leaves, whereas trees occurring within the blue frame might grow successfully if they had more branches. Feedback, where available (see below) was presented for a period of 500ms (800ms for missed trials) and consisted of a numerical reward (if the tree grew successfully) or penalty (if it did not) for planting a tree, and always a reward of zero for not planting a tree. At the beginning of the feedback period, the tree stimulus was replaced by a fixation cross and the response contingencies were replaced by numeral rewards. These rewards/penalties were mapped onto the relevant dimension (branchiness/leafiness) and hence varied in five discrete steps from -50 to +50. Rewards (values above 0) were displayed in green, whereas penalties (rewards below zero) were displayed in red. Rewards of zero were displayed in black. Again, the chosen option was highlighted by a rectangle, with its colour matching the colour of the reward value (red/green/black). For training sessions, the intertrial interval (ITI) had a duration of 1000ms. The directionality of the rewards (more vs less leafy/branchy trees grow better) and the task order during the main training phase were fully counterbalanced across participants.

The training session consisted of three different blocks in which contexts could be either *blocked* or *interleaved. Blocked* means that all trials of in one context were presented first, followed by all trials in another context, with the order counterbalanced over participants. *Interleaved* means that trials were shuffled so that they occurred in random order, but with exactly the same number in each context. Participants underwent a brief interleaved familiarisation phase with feedback (50 trials), followed by an interleaved baseline test (200 trials, no feedback). There was then a long main training session which was blocked (900 trials) (**Fig. S1a**). The purpose of the baseline training and test was to familiarise the subjects with the task and to assess the effectiveness of the main training.

#### Scanning Session (Day3)

The test session consisted of a brief refresher phase (interleaved, 50 trials, feedback) and the main test phase (interleaved, 600 trials, no feedback). The refresher was completed on the experimenter’s laptop and was identical to the baseline training on day 2. For the test phase inside the scanner, we used a jittered ITI of 2000-6000ms (uniform) during which only the grey background was displayed. The total length of all ITIs was restricted such that all runs had equal length.

#### Psychophysical Model of human choices

To quantify sources of error in the choice patterns, we fit a psychophysical model to the choices of each participant. The model assumed that each tree was categorised with respect to a linear category boundary in tree space, via a logistic choice function. The model comprised four free parameters: (1) angle of the decision boundary in tree space (the boundary was assumed to always pass through the centre of the 2D space), (2) a decision bias or offset to the inflection point of the logistic function; (3) the slope of the logistic function (iv) a proportion of random lapses. The model is identical to that in ref [37] where it is described in more detail. From the estimated category boundary, we calculated an angular bias, quantifying the absolute disparity between the estimated and ground-truth task-specific category boundaries. The model was fitted to human choice by minimising the difference between empirical and predicted choice patterns.

#### Group level inference

For all human analyses, group-level inference was performed via paired t-tests on accuracies and signed-rank tests on parameter estimates. To calculate effect sizes, we report Cohen’s d and its nonparametric equivalent Z/sqrt(N).

#### fMRI Acquisition

Magnetic resonance images were recorded with a 3T Siemens scanner with a 32-channel head coil. A high-resolution T1-weighted structural image (voxel size = 1×1×1 mm, 176×256×256 grid, TR=1900ms, TE=2.52ms, TI=900ms) was acquired for each participant prior to the task. Each fMRI image contained 32 axial echo-planar images (EPI) in descending sequence (3.5×3.5×3.5mm isotropic, slice spacing 4.2mm, TR= 2000ms, flip angle = 80, TE = 30ms). We collected fMRI data in six independent runs of 345volumes each.

#### fMRI Pre-processing

Pre-processing was conducted in MATLAB with SPM12 and custom scripts. For each participant, functional scans were first realigned to the first scan. As EPIs were acquired in descending sequence, we applied a slice time acquisition correction with the middle slice (TR/2=1s) as reference. Next, the structural scan was co-registered to the mean EPI. Anatomical scans were normalised to standard Montreal Neurological Institute (MNI) 152 template. EPIs were normalised to the template using tissue probability maps for grey matter, white matter, and cerebrospinal fluid. The EPIs were resliced to 3×3×3mm resolution. For univariate analyses, we applied smoothing with a full width half maximum (FWHM) Gaussian kernel of 8mm.

#### fMRI Data Analysis: GLMs

Data were analysed using SPM12, the RSA toolbox **[49]** and custom scripts written in MATLAB. We used a general linear model (GLM) approach for all univariate analyses. A 128s temporal high-pass filter was applied to remove low-frequency scanner artefacts. Temporal autocorrelation was estimated with a first-order autoregressive model (AR-1). All GLMs contained regressors coding for onset and duration (boxcar until participant response) of events, which were convolved with the canonical haemodynamic response function (HRF). Six motion parameter estimates from the pre-processing stage were included as nuisance regressors in all GLMs. Each run was represented by a separate set of regressors in the GLM, and run number was encoded by a dummy variable. Observed fMRI data at single subject level was regressed against this design matrix. Our analyses are based on three different GLMs. The first GLM (GLM1) had two predictors of interest (task switch trials and task stay trials), locked to cue onset. GLM2 included two parametric regressors of absolute distance of stimuli to the category boundary, for the relevant and irrelevant dimension, respectively. GLM3 was constructed for representational similarity analysis (RSA) and fitted to unsmoothed EPIs. It had 50 regressors per run, one for each combination of context (“north garden”/blue rectangle vs “south garden”/orange rectangle), branchiness (1 to 5) and leafiness (1 to 5).

#### Representational Similarity Analysis of human fMRI

GLM3 (described above) was fit to neural data at single-voxel level. We then constructed neural Representational Dissimilarity Matrices (RDMs) using a spherical searchlight (radius 12mm). For each searchlight sphere, we computed cross-validated neural RDMs from the condition-by-voxel matrix of estimated neural responses using Pearson correlation distance between pairs of conditions from distinct runs. This yielded a 300×300 RDM (50 conditions per run, six runs). All analyses excluded within-run similarity data (e.g. blocks of 50 conditions on the major diagonal). We constructed seven model RDMs to probe for the existence of task-related representational geometries in the fMRI activity patterns: the (1) grid model, (2) orthogonal manifold model, (3) parallel manifold model and (4) rotated grid model, (5) only branchiness model, (6) only leafiness model and (7) diagonal model. The first model encoded two parallel, evenly spaced grids (unit distance), representing each combination of context, branchiness and leafiness. The second model was obtained by taking the grid model and projecting stimuli onto the task-relevant axes for each context. Thus, for each context, stimuli differed along the task-relevant dimension (unit distance), and representations of different tasks were orthogonal to each other. The third model was obtained by rotating one of the task vectors from the second model by 90 degrees, considering the reward assignment the participant had been trained on (hence discriminating “plantiness” of trees, i.e. the extent to which “plant” was the correct answer). For the fourth model, we performed the same rotation on the grid model. The fifth and sixth models served as controls, based on the assumption that early visual areas might exhibit task-agnostic shape (branchiness) or colour (leafiness) sensitivity. The last model was obtained by taking the grid model and projecting trees onto the main diagonal, ranging from low leafiness and low branchiness to high leafiness/branchiness. This was based on the competing hypothesis that humans may have ignored context and optimised for a strategy that yielded 70% correct on both tasks **[37]**. Within a given structural ROI or searchlight sphere, z-scored neural RDMs were regressed against z-scored sets of model RDMs using a multiple linear regression at single subject level. Statistical inference was performed with a group-level t-test of the regression weights against zero. Correction for multiple comparisons was conducted via non-parametric cluster correction as implemented in the SNPM toolbox (FDR threshold < 0.05). To avoid circular inference, all post-hoc visualisations and analyses within ROIs were performed in leave-one-subject-out cross-validated ROIs derived from the activity peaks identified with the searchlight approach (12 mm radius).

#### fMRI RSA: Parametrised Model

In order to obtain more fine-grained estimates of the neural geometry, we also fit a parametrised model to the cross-validated ROIs identified with the searchlight approach. We constructed a space of model RDMs by varying six parameters, one controlling the angle between the task-specific grids (ranging from parallel over orthogonal to anti-parallel in steps of 1degree), four controlling for the compression of relevant and irrelevant dimensions within each context, and one controlling for the separation of contexts. We fit RDMs derived from this model to neural RDMs using a constrained optimisation procedure (fmincon in MATLAB) with least-squares cost function. We then performed group-level inference on the distribution of best-fitting parameter values. These were used to visualise the representational geometries of the best fitting RDMs via projection into three dimensions with classical Multi-Dimensional Scaling (MDS).

#### fMRI RSA: Intrinsic Dimensionality

We performed Singular Value Decomposition (SVD) on the patterns of BOLD activity across voxels within each cross-validated ROI and calculated the cumulative explained variance based on the squared singular values to obtain an estimate of the intrinsic dimensionality of the neural activity patterns. To test whether the directions of largest variance were aligned with the task-diagnostic dimensions of context, branchiness and leafiness, we repeated the regression-based RSA within each cross-validated candidate region after successively removing components, starting with the smallest one. This truncated SVD allowed us to identify the minimal number of components required to successfully decode a factorised representation from the neural data.

#### fMRI RSA: Correlations between brain and behaviour

We performed a correlation analysis (Kendall’s tau) to quantify the extent to which orthogonal representations at the neural level predicted accurate, axis-aligned behavioural responses. We analysed human choice patterns by computing behavioural data RDMs from the probabilities of responding “plant” to trees in each condition, i.e. as a function of each stimulus’ distance to bound along the irrelevant and relevant dimension in each context. Building on previous work **[37]** we fit two model RDMs to human choice patterns, called the *factorised* and *linear* models. In the factorised model, choices were aligned with the ground-truth boundaries, whereas in the linear model, a “diagonal” boundary was applied to both contexts, corresponding to the single linear boundary that optimised for accuracy whilst ignoring the context (yielding ∼70% correct). Fitting the factorised model to behaviour yielded an “axis-alignment score”, indicating whether the participant’s decision boundaries were aligned with the ground truth. We tested at the group level whether the extent to which neural geometries could be explained by the orthogonal model (neural factorisation score) significantly covaried with the extent to which the factorised model explained human choices (axis alignment score).

#### Neural Network Simulations

All neural network simulations were implemented and analysed in Python using the NumPy, SciPy and Scikit-Learn packages. Due to the simplicity the architecture, gradients and optimisation procedures could be derived by hand and implemented in raw NumPy.

#### Task Design

For all neural network simulations, we replaced the fractal tree images with two-dimensional isotropic Gaussian “blobs”. The stimulus space was spanned by parametric modulation of the x and y coordinates of these blobs in five discrete steps. Inside this 5×5 grid, neighbouring blobs were partially overlapping, allowing the network to infer similarity structure based on co-activation of input units. We used a similar context-dependent decision-making task as for our human participants. There were two contexts, in each of which only one feature dimension (either the x- or y-location) was diagnostic of the correct output (the other being an irrelevant dimension) and mapped onto a numerical reward ranging from -2 to 2. The network was trained to predict the reward received in each situation. To assess performance and representational geometries, we fed trials covering all combinations of the two feature dimensions (x/y location) and context into the network and recorded hidden layer activity patterns as well as network outputs for each stimulus.

#### Neural Network Architecture

Our model was a feed-forward network architecture with a single hidden layer. Input units encoded pixel intensities of vectorised and normalised images of Gaussian blobs. Each image had a down-sampled resolution of 5×5 pixels, hence resulting in 25 stimulus input units. Two additional one-hot encoded inputs (1 or 0) signalled the context to the network. All 27 inputs were projected into a hidden layer with 100 units, which were in turn passed through Rectified Linear Unit (ReLU) nonlinearities. The hidden units projected onto a single linear output unit.

#### Weight Initialisation

All network parameters were initialised with random draws from Gaussian distributions with a mean of zero. To control whether the network operated in the rich or lazy regime, we modified the variance of these distributions systematically, ranging from 0.01 (rich regime) to 3 (lazy regime). We call this “initial weight scale” in the main text. These values were derived empirically by observing their impact on the relative change of the weight norm and shape of the loss trajectories during training. Weights to the output unit were instead initialised with a variance scale of 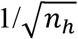 where *n*_*h*_ is the number of hidden units. All biases were initialised to zero.

#### Training

We collected 30 independent runs (unique random initialisations) per initial weight scale condition. On each run, the network was trained with minibatch gradient descent (batch size 50, interleaved data, learning rate 0.001, SGD optimiser) on 10000 iterations. The model was trained on the Mean-Squared-Error (MSE-Loss) between the true and predicted reward associated with each stimulus.

#### Addition of Gaussian Input Noise

We investigated the robustness of different training regimes to additive Gaussian noise in the inputs. The model architecture and training procedures were identical to the ones described above. Again, we collected 30 independent runs per weight scale, ranging from 0.01 to 3 in eight steps. However, this time, we added Gaussian noise drawn from a standard normal distribution to the input units at test. The strength of this noise was varied parametrically in 10 steps from 0 to 0.1, allowing us to investigate the impact of different noise levels on performance.

#### Endpoint Weight Norm and Relative Weight Change

Every 100 epochs during training, we computed the Frobenius norm of the hidden layer weights and their relative change with respect to the norm at initialisation. This allowed us to assess whether the network operated in the rich or lazy regime, corresponding to low and high norm solutions. The weight change relative to initialisation was quantified by computing how the norm of the hidden layer weights changed from random initialisation to the endpoint of training.

#### Neural Network Representational Similarity Analysis

We performed RSA on the hidden layer activity patterns to assess how training sculpted the representations formed by the neural network. For each individual run, we calculated RDMs based on the hidden layer activity patterns evoked by inputs covering all combinations of feature values and contexts. The resulting 50×50 RDMs captured the Euclidean distances between all possible pairs of stimuli in the high-dimensional space spanned by the hidden units (after the ReLU nonlinearity). We visualised these geometries by projecting the group-level RDM, averaged across independent runs, down into three dimensions using metric MDS.

#### Neural Network RSA: Quantifying hidden layer geometries

To quantify the extent to which hidden layer geometries exhibited patterns consistent with our hypotheses, we performed a linear regression of the hidden layer RDMs onto a set of model RDMs. There were three model RDMs in total, (1) a grid model, encoding the stimulus spaces as two parallel grids, separated by the context, (2) an orthogonal model, encoding the task relevant dimensions as two orthogonal 1D manifolds and (3) a parallel model, encoding the same information as the orthogonal model, but rotated into the frame of reference of the response (i.e., a “magnitude” representation). The lower triangular form of these models was z-scored and entered into a linear multiple regression model to predict the lower triangular form of the hidden layer RDM. This procedure was repeated for each individual run, yielding a distribution of regression coefficients that permitted statistical inference on the relative difference between predictors as well as their difference from zero. We tested whether two models differed in their extent to which they covaried with the hidden layer RDM by performing Wilcoxon Signed Rank tests on their corresponding beta estimates. A nonparametric test was chosen due to the observed violation of the normality assumption. We applied this analysis to models with different initial weight scale, enabling us to investigate the impact of the training regime (rich or lazy) on the emerging representations.

#### Neural Networks: Intrinsic Dimensionality of hidden layer activity patterns

We used SVD to investigate the dimensionality of the hidden layer activity patterns. SVD was applied to the stimulus-by-unit matrix of hidden layer responses to all combinations of feature values and context. We visualised the cumulative variance explained based on the squared singular values (i.e., the eigenvalues of the response matrix) as Scree plot and performed the Elbow method to obtain a qualitative estimate of the intrinsic dimensionality. Next, we performed truncated SVD to assess the task-diagnosticity of the first ***k*** directions of variation in the response matrix. For this, we reconstructed the hidden layer response matrix, keeping only the ***k*** first singular values with ***k*** ranging from 1 to 27 (i.e. the number of input units). We then generated new outputs from the network by passing this lower-dimensional activity pattern on to the output unit. Lastly, we calculated the accuracy as the mismatch between these outputs and the ground truth. This allowed us to assess, separately for the rich and lazy regime, the extent to which removing components from the hidden layer responses reduced the network’s performance. The hypothesis was that more components would be needed in the lazy compared to the rich regime to maintain equal task accuracy.

#### Neural Network Hidden Unit Selectivity and Axis Alignment

To investigate task selectivity of hidden layer units, we capitalised on the property of ReLU nonlinearities that they map negative inputs to zero. We defined task-selectivity for the neural network as a non-zero response to stimuli in one context and zero response to all stimuli in the other context. Stimulus selectivity irrespective of context was defined as having a non-zero response in both contexts. We calculated these sensitivity indices at initialisation and after training to ensure that the initialisation scheme did not pre-partition the hidden layer in the absence of a training objective. Dead units were defined as returning zero for all stimuli (all combinations of feature values and context). From this, we calculated the proportion of units that were either dead, task- or stimulus-selective. To visualise response profiles, we averaged activity within these sub-populations, constructed a response matrix of these averages separately for each context (with rows corresponding to y location, columns to x-locations of stimuli and the value corresponding to the average activity of a sub-population) and plotted the group level average (mean across independent runs) as heatmaps. For this, we focussed on the two most extreme weight initialisations, 0.01 and 3, corresponding to learning in the rich and lazy regime, respectively. Lastly, to quantify the extent to which these response patterns were axis aligned (i.e., whether units responded to relevant but not irrelevant dimensions), we concatenated the two vectorised task response matrices, constructed RDMs based on pairwise differences in magnitude and regressed them against two model RDMs, (1) the axis-aligned and (2) diagonal models. In the axis aligned model, unit responses scaled with context-dependent relevant dimensions (i.e., with x-location in context A and y-location in context B). In the diagonal model, activity scaled jointly with both dimensions irrespective of context. We fitted the model at the level of individual runs. To assess which model RDM covaried stronger with the observed neural responses, we performed a Wilcoxon Signed Rank test on the difference between beta estimates for the axis-aligned and diagonal model. To assess whether this difference was dependent on the initialisation scheme, we performed the same test on the difference of differences.

#### Neural Network context weight correlations

Our theory predicted that the network could learn the gating scheme via anti-correlated context weights. To test this empirically, we calculated the Pearson correlation between task A and task B weights from the input to the hidden layer at the level of single runs both at initialisation and after the last training epoch. We repeated this analysis on the sub-populations of task-selective and stimulus-selective units, expecting weights into the former to be stronger anti-correlated. We visualised the distribution of single-run correlation coefficients together with a Kernel-Density-Estimate computed with the kdensity function from the Seaborn package.

#### Neural Network Ablation Study

We performed an ablation study to investigate how critical task-sensitive and stimulus-sensitive units were for multi-task performance. More specifically, for each collected run, we set either the sub-population of task-selective or stimulus-selective units to zero, performed a forward pass through the ablated network and computed its loss and accuracy.

#### Nonhuman primate data

NHP results were based on a reanalysis of data recorded from monkey frontal eye fields (FEF) during performance of comparable context-based decision-making tasks. The data are freely available at https://www.ini.uzh.ch/en/research/groups/mante/data.html. These data have already been intensively scrutinised in past work [4,36]. In the experiment, two monkeys were asked to discriminate between distinct levels of motion direction and colour of random dot stimuli, with only one dimension being relevant in each context, just as in our experiments. Stimuli spanned a similar 2D grid (motion directions varying from left to right, colour gradient from green to red) as our trees and Gaussian blobs. Further details are available in ref [4].

#### Representational Similarity Analysis of NHP electrode recordings

We created pseudo-populations by concatenating all recorded units, separately for monkey A and monkey F. Unit-by-stimulus response matrices were obtained by averaging activity across trials for each stimulus type (6 motion directions * 6 colours * 2 contexts = 72 entries). RDMs were constructed from these matrices using the Euclidean distance measure. For all reported analyses, we focus on activity averaged over the second half of the trial (300-600ms) as task factorisation was strongest in this interval, an observation consistent with previous reports of dynamic encoding of different task variables throughout a trial **[36]**. We fitted the same set of candidate model RDMs to this dataset as previously to RDMs obtained from human fMRI data (see above). For statistical inference, we created a null distribution by randomly permuting the trial labels and repeating this regression-based RSA 1000 times. Significance was defined as regression weights two standard deviations above this null.

#### Individual Unit Selectivity and Axis Alignment of NHP electrode recordings

We assessed task selectivity of individual units using a standard regression-based approach. Mean activity of each unit was regressed against four predictors, coding for colour and motion direction separately for each context. Selectivity was defined as having a significant regression coefficient for the variable of interest. Due to the substantial number of tests, we performed FDR correction to correct for multiple comparisons. We distinguish between diverse types of selectivity. Task selectivity was defined as having a significant regression weight only for the relevant feature dimension (i.e., only for motion in the motion task and colour in the colour task). Stimulus selectivity was defined as having significant coefficients for both dimensions. Furthermore, we identified units that were selective only to colour or motion, irrespective of context, and defined “mixed”/non-specific selectivity as having significant regression weights that do not fall into any of the above categories. As for the hidden units in the neural network, we again plotted the different proportions of selectivity patterns of units within a pseudopopulation and visualised the response profile of task and stimulus selective units by averaging the activity within a sub-population separately for each combination of feature values (colour, motion) and context. Axis alignment of these response matrices was assessed by regressing them against the factorised and diagonal model as previously described for the neural network (see above). We assessed the intrinsic dimensionality of the patterns observed in monkey FEF using the same truncated SVD approach described above for the human fMRI data.

## Supplementary Information

Supplementary figures S1 to S8

**Supplementary Figure S1.**
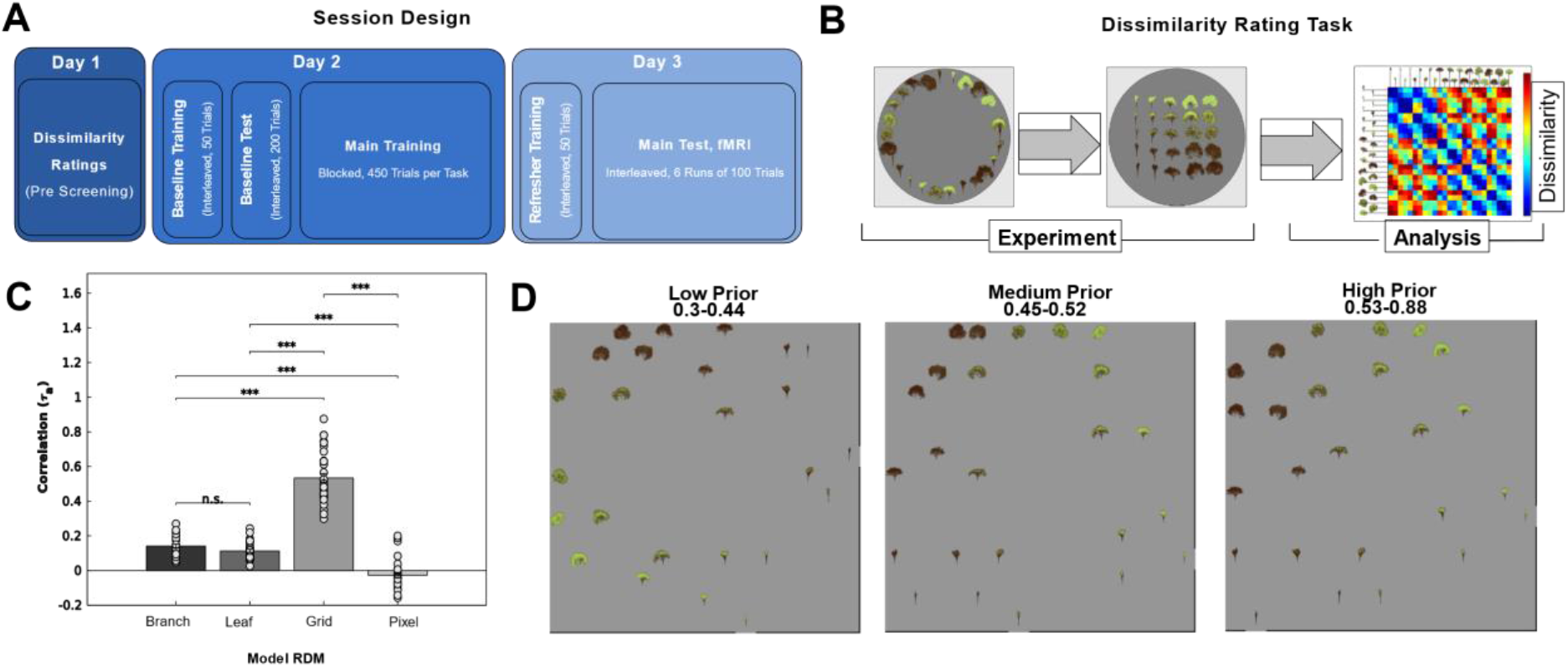
**(a)** Session Design. Participants completed three sessions carried out over consecutive days. All participants underwent a screening task (day1) in which they were asked to perform dissimilarity ratings on tree stimuli. Those who showed strong evidence for being aware of the dimensions of branchiness and leafiness (assessed by a “grid score”, see next figure) were invited to the remaining parts of the study. On day 2, participants received a lengthy blocked training curriculum, preceded by a brief familiarisation phase and evaluation (baseline training and test) to measure the effectiveness of the training phase. On day 3, participants received a brief refresher training, before they underwent fMRI scanning during which they completed six interleaved blocks of test trials. See methods sections for additional details. **(b)** Dissimilarity Rating Task & RSA. Participants were asked to arrange tree stimuli via mouse drag & drop in a circular arena such that distances between trees corresponded to how dissimilar they were perceived (left and middle panel). From these ratings, we constructed RDMs at single subject level. These RDMs were correlated with model RDMs assuming that participants were only aware of branchiness, (ii) only aware of leafiness, (iii) aware of the full 5×5 grid of branchiness and leafiness or (iv) made judgements based on pixel similarity. We describe the extent to which the third model explains the data as “grid score”. In Flesch et al, 2018, we reported interactions between training effectiveness and grid score. We thus only invited participants with a grid score higher than the median grid score (tau=0.18) from the previous study. All screened participants exceeded this threshold. **(c)** Correlation coefficients between subject ratings and model RDMs. The grid model explained the data best, indicating that participants were on average aware of the data-generating dimensions. **(d)** MDS on dissimilarity ratings, divided into participants with low, medium and high grid score. All groups showed evidence for awareness of the dimensions branchiness & leafiness, and their grid-like relationship with each other.

**Supplementary Figure S2.**
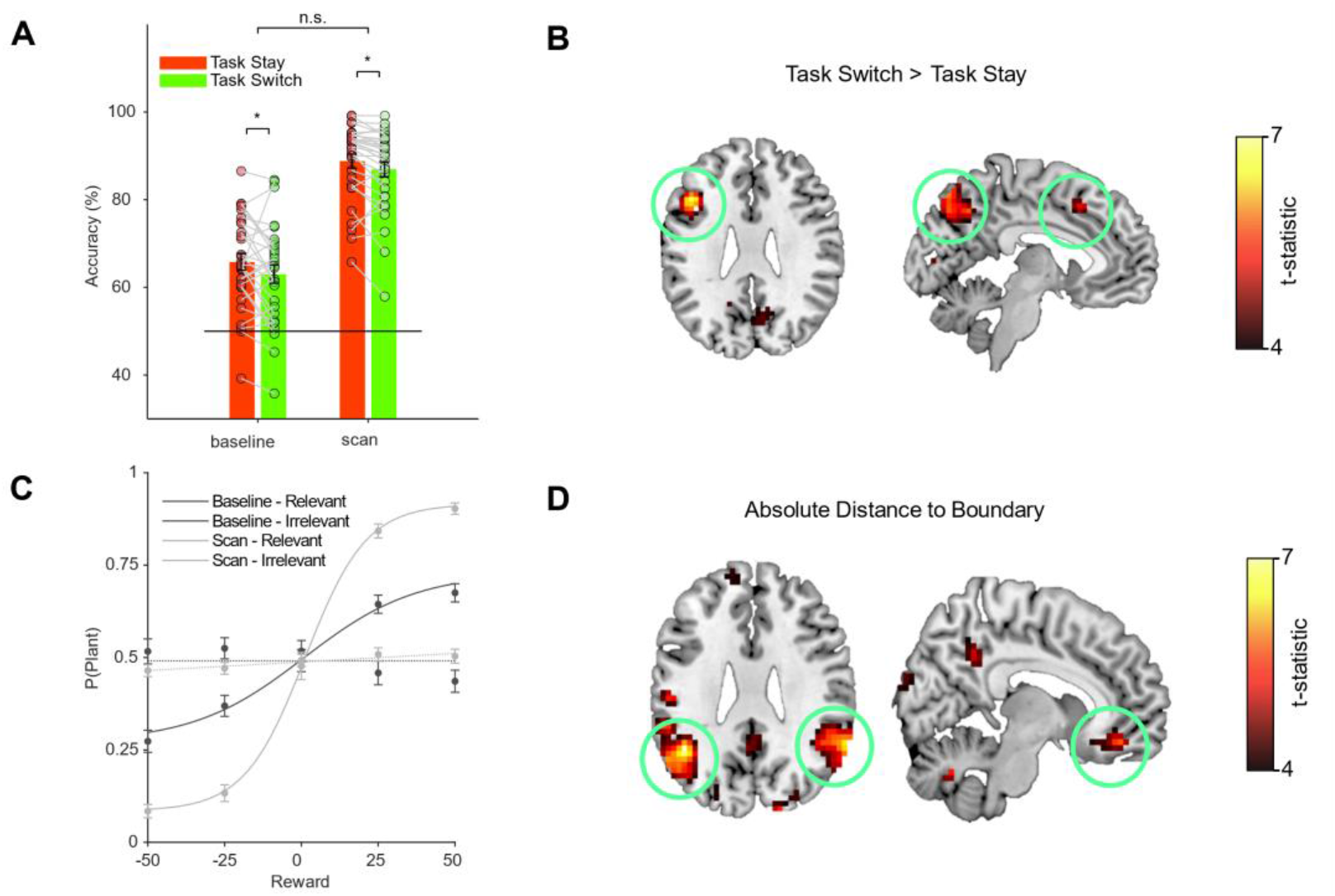
**(a)** Behavioural switch cost. Participants were slightly worse on switch than stay trials at test, both during the baseline and later scanning session (*Accuracy Baseline, Switch < Stay: T(29)=2*.*057, p=0*.*048, d=0*.*266; Accuracy Scan, Switch < Stay: T(29)=2*.*715, p=0*.*011, d=0*.*211; Interaction Phase x Switch cost: T(29)=-0*.*668, p=0*.*509, d=-0*.*251*). **(b)** Univariate markers of switch cost. A whole-brain univariate contrast of switch vs stay trials revealed lusters in task-positive regions where activity was higher on switch than on stay trials. More specifically, we found significant clusters in Parietal Cortex *(BA7* : *t(30) = 5*.*65, p < 0*.*001 (FWE corrected), cluster extent (kE) = 570, MNI coords = [-6, -74, 52])*, Supplementary Motor *Area (SMA t(30) = 5*.*03, p < 0*.*05, kE =66, [-6, 18, 46]))* and left Medial Frontal Gyrus *(MFG t(30)=6*.*55, p<0*.*01, kE = 124, [-44, 21, 28]))* **(c)** Behavioural sensitivity to relevant and irrelevant dimensions. Fitting logistic functions to the choice patterns along both dimensions revealed that, compared to the baseline, participants became much more sensitive to the task-relevant dimension after they had engaged in the blocked training phase *(Slope Relevant, Baseline: Z = 4*.*72, p = < 0*.*001, d = 0*.*873; Scan > Baseline: Z = 4*.*762, p = < Scan: Z = 2*.*705, p = 0*.*007, d = 0*.*494)*. Participants were, however, much more sensitive to the relevant than irrelevant dimension at *test (Scan, Relevant > Irrelevant: Z = 4*.*782, p < 0*.*001, d = 0*.*873)*, and this sensitivity was higher compared to baseline *(Dimension x Phase Interaction: Z = 4*.*741, p < 0*.*001, d = 0*.*866)* **(d)** Univariate markers of absolute distance to category boundary. A GLM with parametric regressors for the absolute distance to category boundary (methods) revealed significant relationships between activity and distance to bound along the relevant, but not irrelevant feature dimensions. More specifically, we found significant clusters in bilateral Angular Gyrus *(left: t(30) = 6*.*79, p < 0*.*001, kE = 364, [60, -49, 28])* and the right Orbitofrontal Corex (t(30) = 5.46, p < 0.01, kE = 73, [8, 42, -14]), and to a lesser extent also in bilateral EVC *(left: t(30) = 5*.*15, p < 0*.*01, kE = 70, [-13, -98, 14]; right: t(30) = 6*.*55, p < 0*.*01, kE = 61, [18, -94, 21]*) as well as the Posterior Cingulate cortex *(t(30) = 5*.*05, p < 0*.*001, kE = 192, [4, -49, 35])*.

**Supplementary Figure S3:**
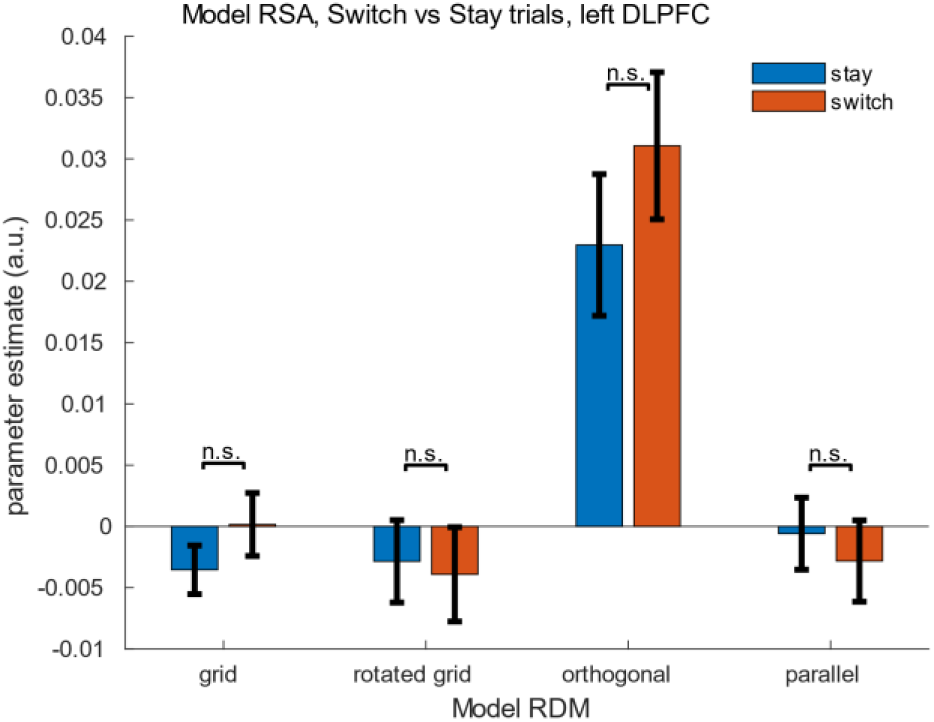
Model RSA separately for switch vs stay trials. The univariate contrast of switch vs stay trials revealed a significant difference in BOLD in left DLPFC, an area where we had also observed evidence for factorised representations using the searchlight RSA approach. We therefore tested whether the extent to which task representations were factorised (i.e. lied on orthogonal manifolds) differed between switch and stay trials. The difference, however, was not significant.

**Supplementary Figure S4.**
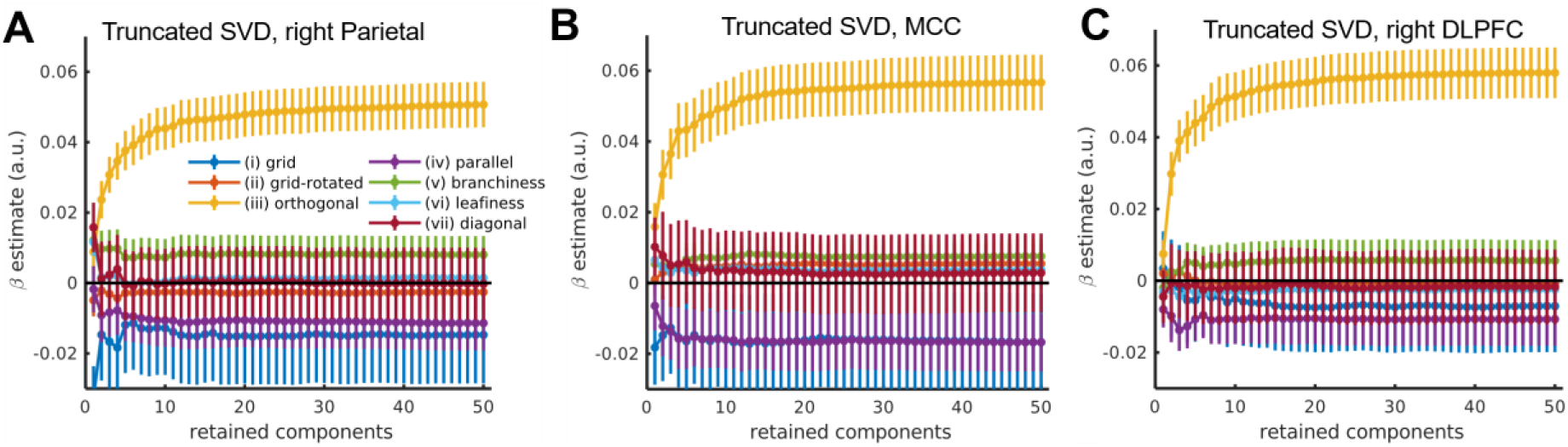
Truncated SVD on fMRI patterns. To estimate the intrinsic dimensionality of the neural activity manifold, we repeated the model-based RSA on reconstructions of the data for which we had successively removed principal components (truncated SVD, see methods). Across all three regions, we observed that orthogonal representations could be reliably decoded if only the two strongest components were retained, and there was no measurable benefit in retaining more than the six largest PCs. Together, these results indicate that the largest directions of variance are to some extent aligned with the directions spanning the orthogonal task manifold, indicating that this manifold is intrinsically low-dimensional.

**Supplementary Figure S5.**
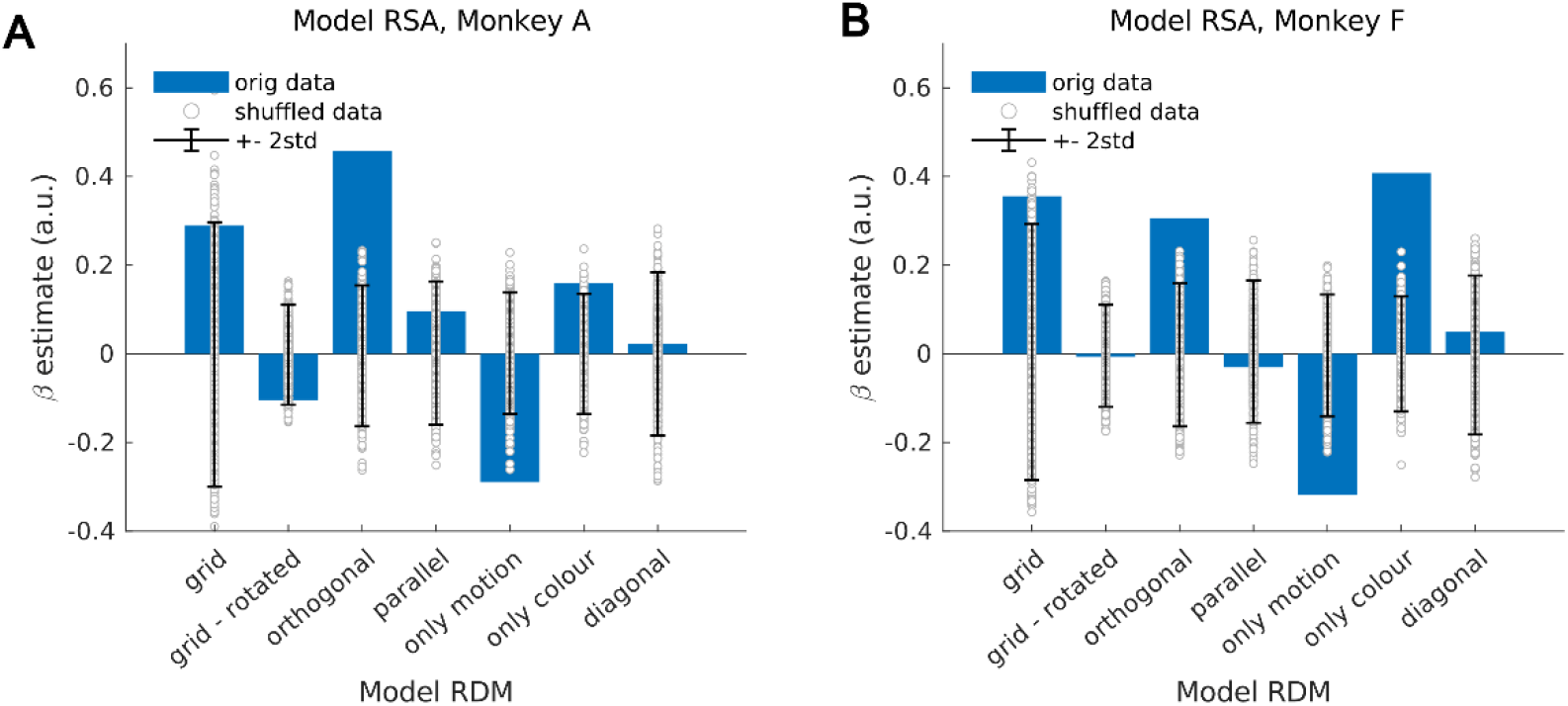
Model-based RSA on NHP data (both monkeys). We fitted the same set of model RDMs to the monkey data as to our human participants (see methods). We found strong evidence for orthogonal representations, encoding only relevant feature dimensions, in monkey A. In contrast, neurons recorded from Monkey F responded predominantly to colour, irrespective of the task the monkey was doing, which is consistent with previous reports of heterogenous responses in the two monkeys (Mante et al, 2013)

**Supplementary Figure S6.**
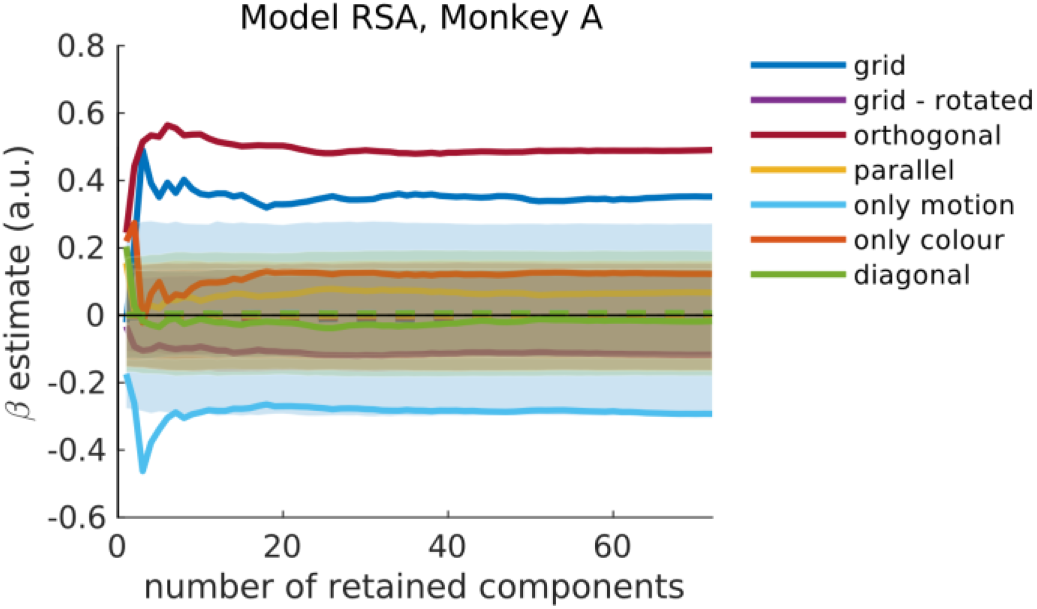
Truncated SVD on NHP data. We subjected the neural RDMs from Monkey A to the same truncated SVD approach as the RDMs from human fMRI data, to assess the intrinsic dimensionality of the manifold that encodes task relevant and supresses task irrelevant dimensions. Consistent with our observation in humans, we found evidence for a low-dimensional manifold, as retaining the first two components was sufficient to decode orthogonal representations, which suggests that the strongest directions of variance are aligned with the dimensions spanned by an orthogonal & low-dimensional task manifold.

**Supplementary Figure S7:**
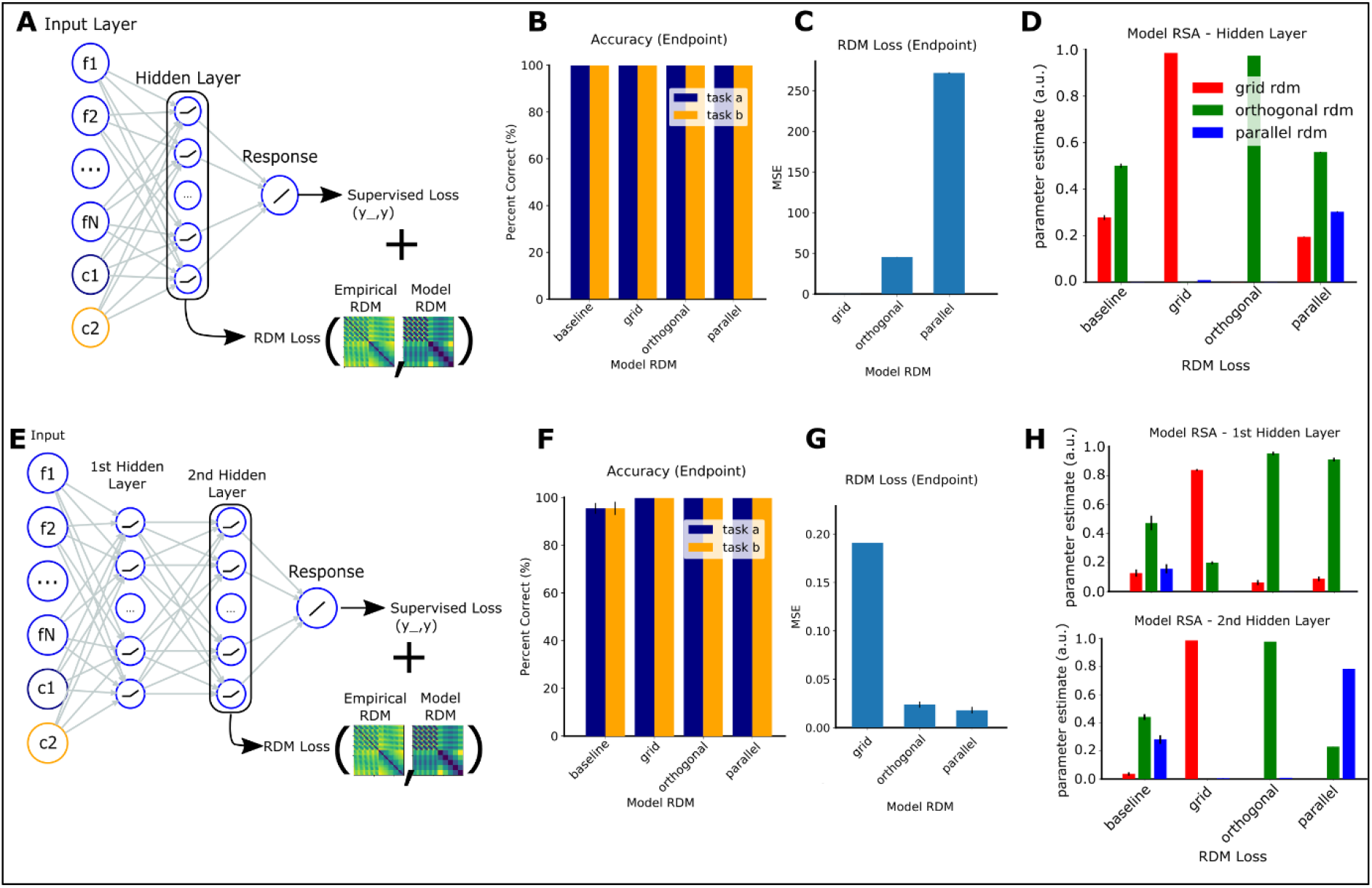
Enforcing specific representational geometries through auxiliary loss function under rich learning. We equipped the network with an auxiliary objective (“RDM loss”) which minimised the difference between patterns in the hidden layer and a candidate model RDM that encoded either grid-like, orthogonal or parallel representational schemes. (A) Illustration of experimental set-up for model with a single hidden layer. (B) Accuracy after convergence on the supervised objective, as a function of the model RDM used for the RDM loss function. All models converged. (C) Endpoint RDM loss after convergence on the supervised objective. The network for which we tried to enforce parallel representations failed to minimise the RDM loss, all other models converged. (D) Regression coefficients for model RSA as a function of the model used for the RDM loss. Baseline corresponds to the vanilla network without an RDM loss. The models with grid and orthogonal schemes as target for the RDM loss learned the desired representations. The model trained with a parallel RDM as target in the RDM loss converged to orthogonal representations. (E) Same as A, but for model with two hidden layers. (F) same as B, but for model illustrated in E. (G) Same as C but for model illustrated in E. This time, the RDM loss with a parallel model converged. (H) Same as D, but for model shown in E. With two hidden layers, promoting parallel representations in the second hidden layer through an RDM loss worked, and led to emergence of orthogonal representations in first hidden layer.

**Supplementary Figure S8.**
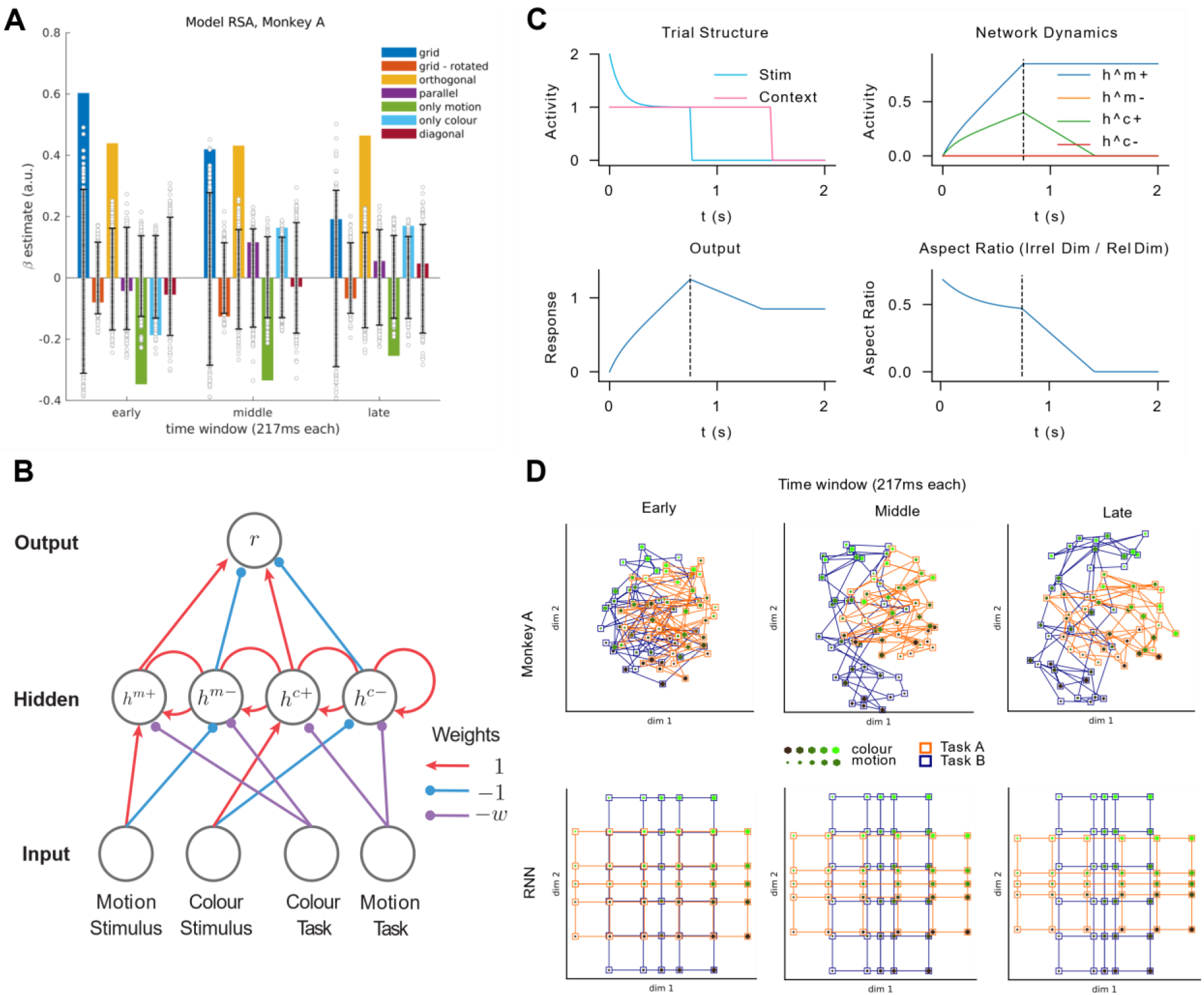
Temporal evolution of orthogonal representations. (A) Model RSA on NHP data, separately for early, middle and late time windows within the stimulus interval. Early in the trial, the grid-model explains the data best. In following intervals, the parameter estimate for the orthogonal model increases progressively, suggesting that the neural code transforms from a grid-like to an orthogonal and task-specific representational scheme. (B) An extension of our network architecture with recurrent hidden connections. Please see supplementary methods for details. (C) Dynamics of RNN throughout a simulated trial. Top left: Stimulus and context signal are presented for 750ms, followed by a delay of 1s during which only context information is provided to the network. Top right: Dynamics of hidden layer activity throughout an example trial where motion direction was relevant. During stimulus presentation, we observe a gradual integration of motion information in the motion-sensitive unit, and, to a lesser extent, colour information in the colour-sensitive unit. After stimulus offset (dashed line), the irrelevant dimension (colour) is gradually supressed by the context signal. Bottom left: Gradual integration of a category signal in the output unit, which remains roughly constant after stimulus offset. Bottom right: Aspect ratio between activity encoding the irrelevant and relevant dimensions respectively, indexing the amount of compression along irrelevant dimensions. The aspect ratio decreases during the stimulus interval as irrelevant and relevant feature information are integrated at different rates (top right plot). It decreases more rapidly after stimulus offset (dashed line) as the context signal filters out any task-irrelevant information that is still present. (D) MDS on monkey and RNN RDMs averaged over early, middle and late time windows within the stimulus interval. In both cases, we observe evidence for a gradual emergence of task-specific and orthogonal representations (with irrelevant features being suppressed) out of more grid-like representational structures.

## Supplementary Methods

### 1 Recurrent Network Extension

Let *x*_1_(*t*) ∈ [-1, 1] be the signed motion coherence over time in a trial, and *x*_2_(*t*) ∈ [-1, 1] be the signed color level over time, which can be stacked into the column vector input *x*(*t*) = [*x*_1_(*t*) *x*_2_(*t*)]^*T*^. Let *u*(*t*) ∈ *R*^2^ be the task context input encoded as a one hot vector (+1 in the first element for context A, +1 in the second element for context B).

The network contains four neuron classes, and the overall architecture is depicted in Fig. 1. In particular, these comprise a pair selective for positive/negative motion and task, and an pair selective for positive/negative color and task. Each neuron receives stimulus input through the input-to-hidden weights

**Figure 1:**
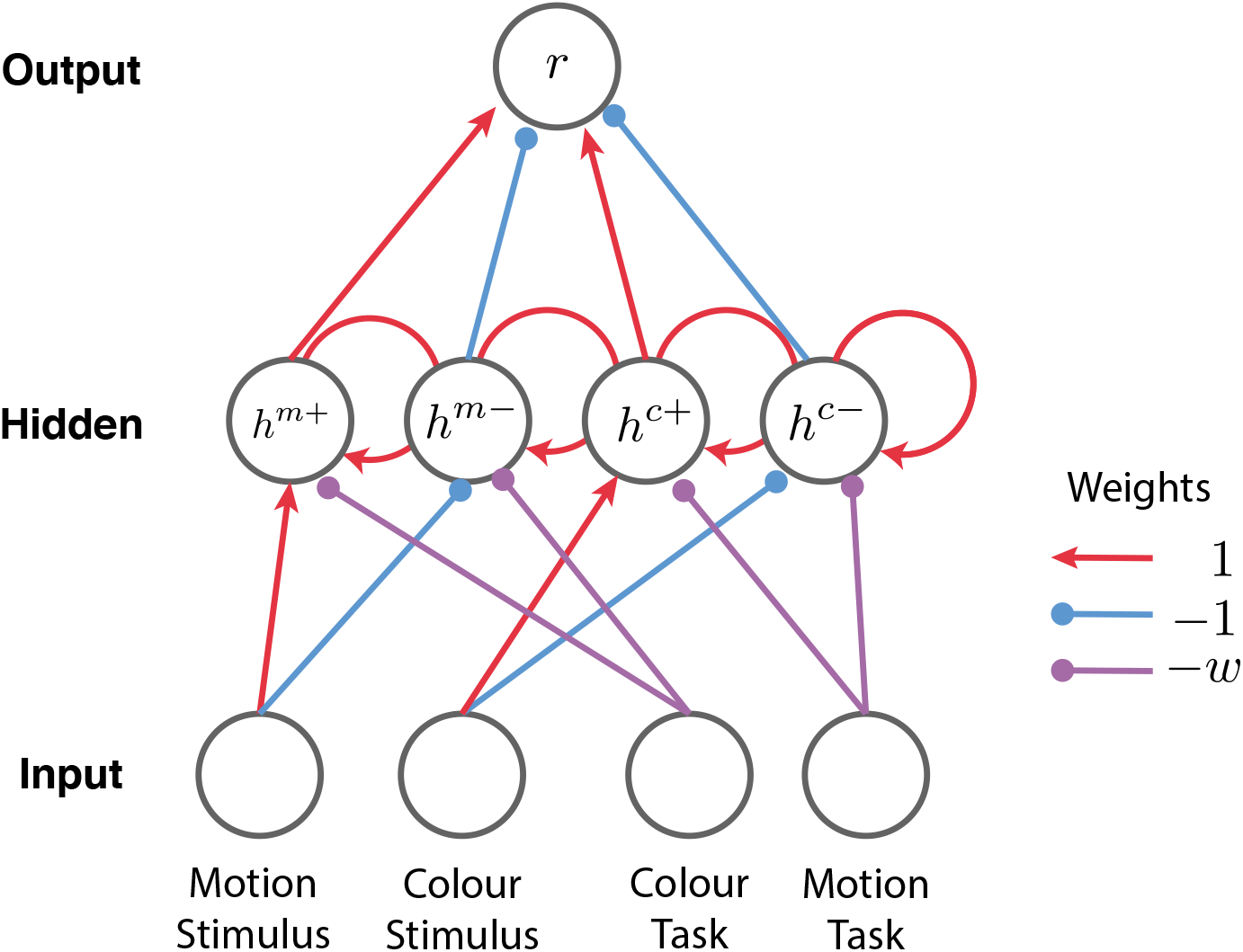
Network architecture for temporal extension.

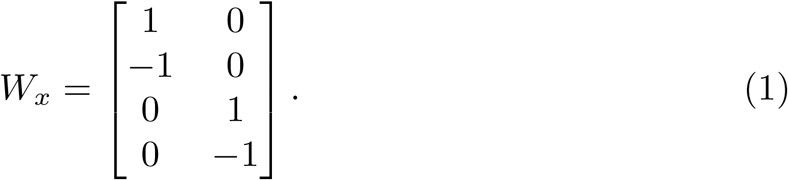

Each neuron class also receives task input, with the motion neurons receiving inhibitory input in the color task and the color neurons receiving inhibitory input in the motion task. The task-to-hidden weights are

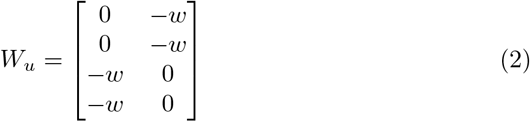

where *w* is a parameter controlling the strength of context-driven inhibition.

The network has recurrence, which we assume has an autapse structure such that each neuron has self recurrence with weight one to enable persistent activity.

We emphasize that all four neuron classes are mixed selective, in the sense that their response depends on a combination of stimulus and task. However, this mixed selectivity is not random, rather it is highly structured.

The neural activity dynamics are given by the standard firing rate equations

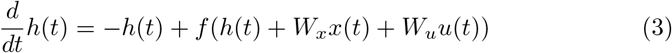

where *f* (·) is the firing rate nonlinearity, which here we take to be the ReLU function (*f* (*v*) = max {*v*, 0}).

Finally the output of the network *r* is computed through readout weights *W*_*o*_ = [1 -1 1 -1], i.e., by summing or subtracting the relevant hidden unit activity,

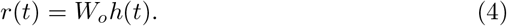

#### 1.1 Input dynamics

We now describe the temporal structure of a trial. We assume that between trials, neural activity resets such that we have the initial condition *h*(0) = 0. We assume that input stimuli arrive with a temporal profile *p*_*x*_(*t*) that is rescaled by the motion coherence *m* and color coherence *c*, such that the input is

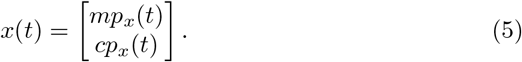

For simplicity we take *p*_*x*_(*t*) = *ae*^*-t/τ*^ +*b* for 0 *< t < t*_*x*_, and *p*_*x*_(*t*) = 0 otherwise, to reflect a sharp onset transient followed by decay to a steady state.

The context signal arrives with a temporal profile *p*_*u*_(*t*), turning on with the stimulus and remaining on during the delay period until some time *t*_*u*_ *> t*_*x*_. For simplicity we take *p*_*u*_(*t*) to be a pulse (one for times between 0 and *t*_*u*_, zero otherwise). Let *z* be 1 in the motion context and 0 in the color context. Then we have

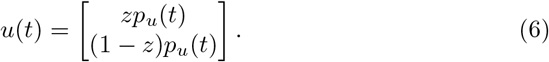

### 2 Solution

The dynamics in this model can be solved exactly. Let 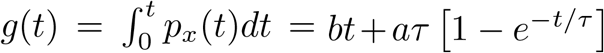 with *a* ≥ 0, *b* ≥ 0, *τ >* 0. Let *h*^*m*+^ denote the positive motion neuron class, *h*^*m-*^ the negative motion class, and so on. We have

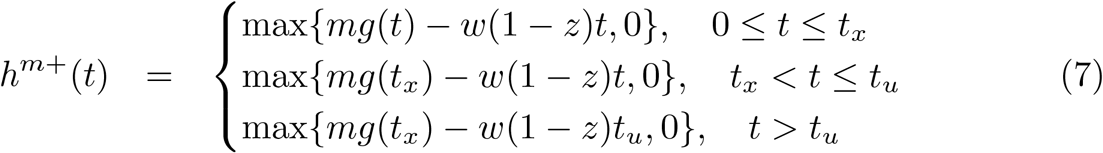

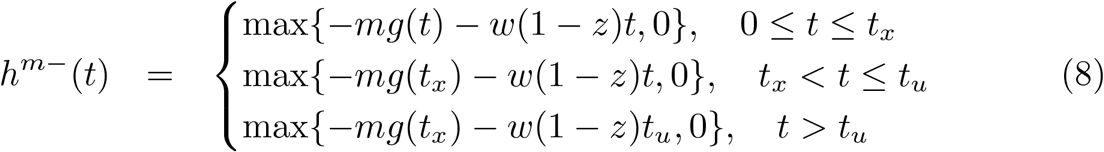

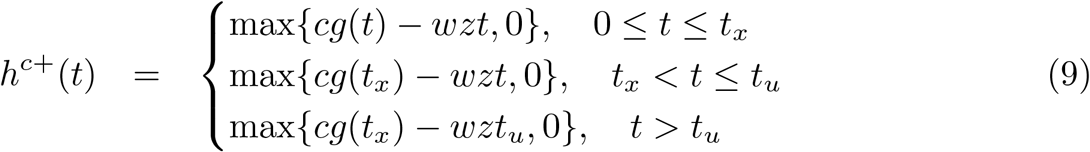

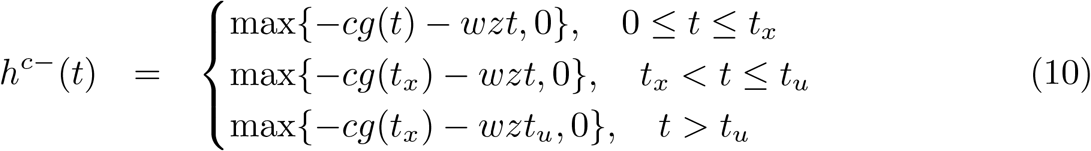

Example dynamics from the model are shown in Fig. 2.

**Figure 2:**
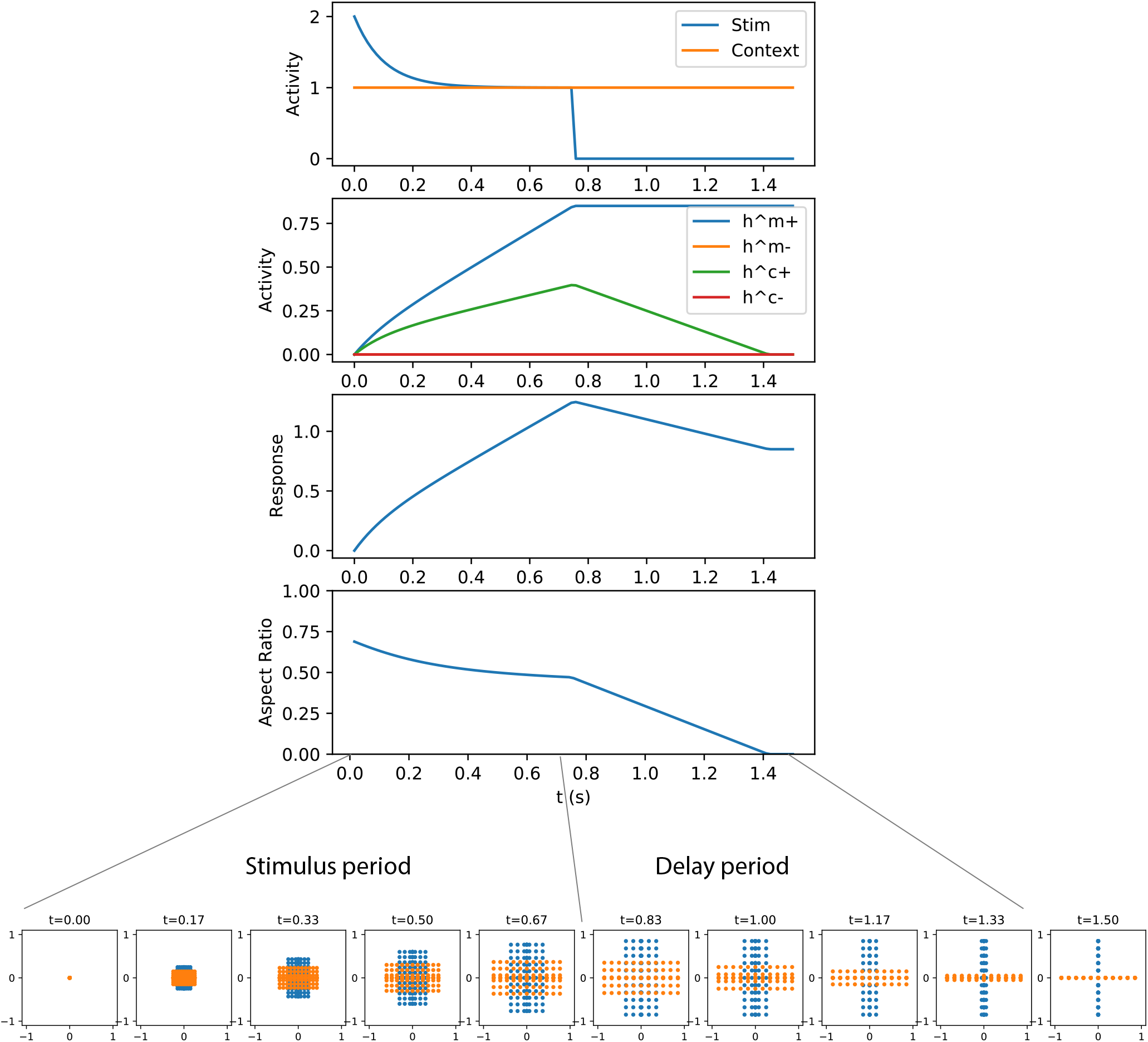
Example dynamics. From top to bottom panel: Input kernels. Activity dynamics for each neuron class. Response variable. Degree of compression of irrelevant stimulus dimension relative to relevant stimulus dimension. Bottom row: 2D representation, showing full grid of motion and color stimuli in each context. Context indicated by color (blue/orange). For these parameters, early time points show little compression compared to later time points. Parameters: *w* = .6, *t*_*x*_ = .75, *t*_*u*_ = 1.5, *a* = 1, *b* = 1, *τ* = 0.1.

#### 2.1 Dimensionality reduction

This activity is four dimensional, but in practice, these dimensions could be combined or rotated in the population response. Common analyses project the population activity down to two or three dimensions using dimensionality reduction techniques. Here we analytically calculate the result of applying Principal Component Analysis (PCA) to perform this reduction. PCA selects the eigenvectors of the hidden activity correlation matrix

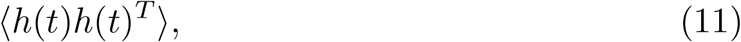

where the average ⟨·⟩ is over the stimulus and task parameters *m, c*, and *z* and the time within trial *t*. Without knowing the details of this average, it is still possible to calculate the principal components. In particular, correlations between positive and negative neuron classes for each stimulus dimension *s* ∈ *m, c* are zero, ⟨*h*^*s*+^*h*^*s-*^⟩ = 0, because only one neuron class is active at a time. Next, correlations across stimulus dimensions will all be equal due to the symmetry in the problem, such that 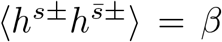 for 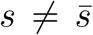. Finally, self correlations will similarly be equal, ⟨*h*^*s±*^*h*^*s±*^⟩ = *α*. We therefore have the correlation matrix structure

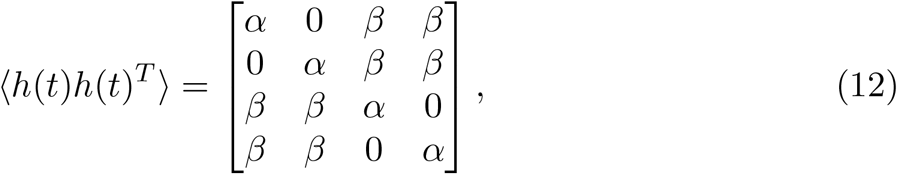

which is ultrametric (consisting of blocks within blocks). All matrices of this form are known to be diagonalized by the Haar wavelets, yielding the orthogonal matrix of eigenvectors,

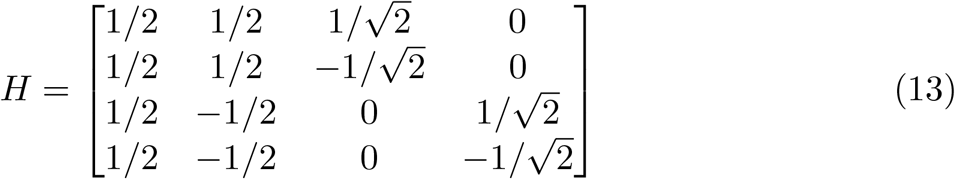

such that ⟨*h*(*t*)*h*(*t*)^*T*^⟩ = *H*Λ*H*^*T*^ where Λ is the diagonal matrix of eigenvalues. These eigenvalues are [*α* + 2*β, α -* 2*β, α, α*]. We can further observe that, for typical settings where irrelevant information is not completely suppressed and motion and colour trials are uniform over a similar range, we expect *α > β >* 0. In this case the largest variance direction is the mean, followed by the two stimulus dimensions, and finally the context offset.

The principal components are the columns of *H* and can be interpreted as the mean activity, the context offset, the motion axis, and the colour axis respectively. In essence, because only one or the other of the ‘positive’ and ‘negative’ populations will be active for each stimulus dimension, PCA forms these into a 2D low dimensional representation *y* where the ‘negative’ neurons are mapped to the negative part of one axis while the ‘positive’ neurons are mapped to the positive part. That is, focusing just on the colour and motion dimensions, we have the transformation

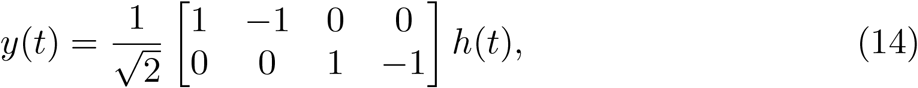

or, for a three-dimensional reduction excluding the mean (as is typical) and including the context offset we have,

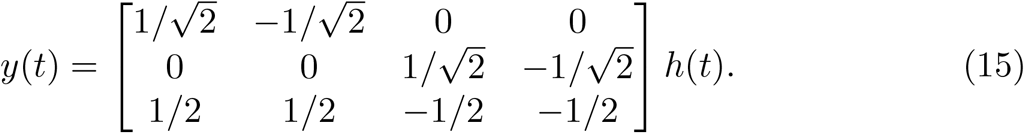

Example dynamics in the 2D space are shown in Fig. 3.

**Figure 3:**
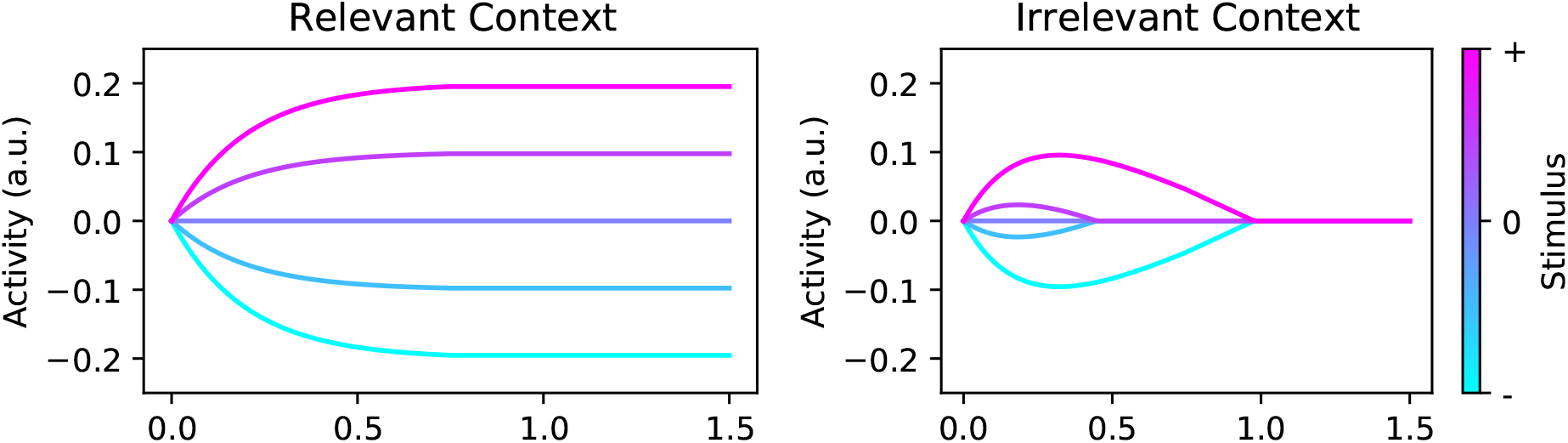
Example lower-dimensional 2D dynamics. Left: Response of low dimensional variable *y*_1_ to several levels of input on stimulus dimension 1 where this stimulus is task-relevant. Right: Response of same variable *y*_1_ to several levels of input on stimulus dimension 1 where this stimulus is task-irrelevant. A brief transient response to stimuli is rapidly suppressed in the irrelevant context. Parameters: *w* = .2, *t*_*x*_ = .75, *t*_*u*_ = 1.5, *a* = 1, *b* = 0, *τ* = 0.2.

#### 2.2 Compression dynamics

To track the changing representation of task-relevant and task-irrelevant stimulus features, we calculate the ratio of the hidden activity for the task irrelevant neuron class compared to the task-relevant neuron class. For instance, if we supply an input *m* = 1, *c* = 1 in the motion context (*z* = 1), then the ratio of the activity of the positive colour neuron class to that of the positive motion neuron class is

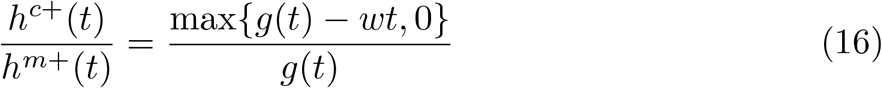

for 0 *< t < t*_*x*_, i.e., during the stimulus period.

### 3 Phase portrait

Finally, we generate a phase portrait for the dynamical system in each context. We construct the phase portrait for *h*^*c*+^ and *h*^*m*+^ and positive *c,m*, with a symmetric situation holding for the other two neuron classes and opposite stimulus sign.

Stimulus input directions are invariant across contexts, with motion stimuli increasing *h*^*m*+^ and colour stimuli increasing *h*^*c*+^. In the motion context, there is a line attractor along the *h*^*m*+^ axis, while in the colour context there is a line attractor along the *h*^*c*+^ axis. In the motion context, activity in the *h*^*c*+^ neuron decays to zero due to the context inhibition, in a direction that is exactly opposite to the colour input direction. Likewise in the color context, activity in the *h*^*m*+^ neuron decays to zero due to context inhibition, in a direction that is exactly opposite to the motion input direction. That is, in each context, the selection vectors’ are orthogonal to the irrelevant input direction. Finally, the readout vector lies at 45 degrees in this space. These features yield the phase portrait shown in Fig. 4.

**Figure 4:**
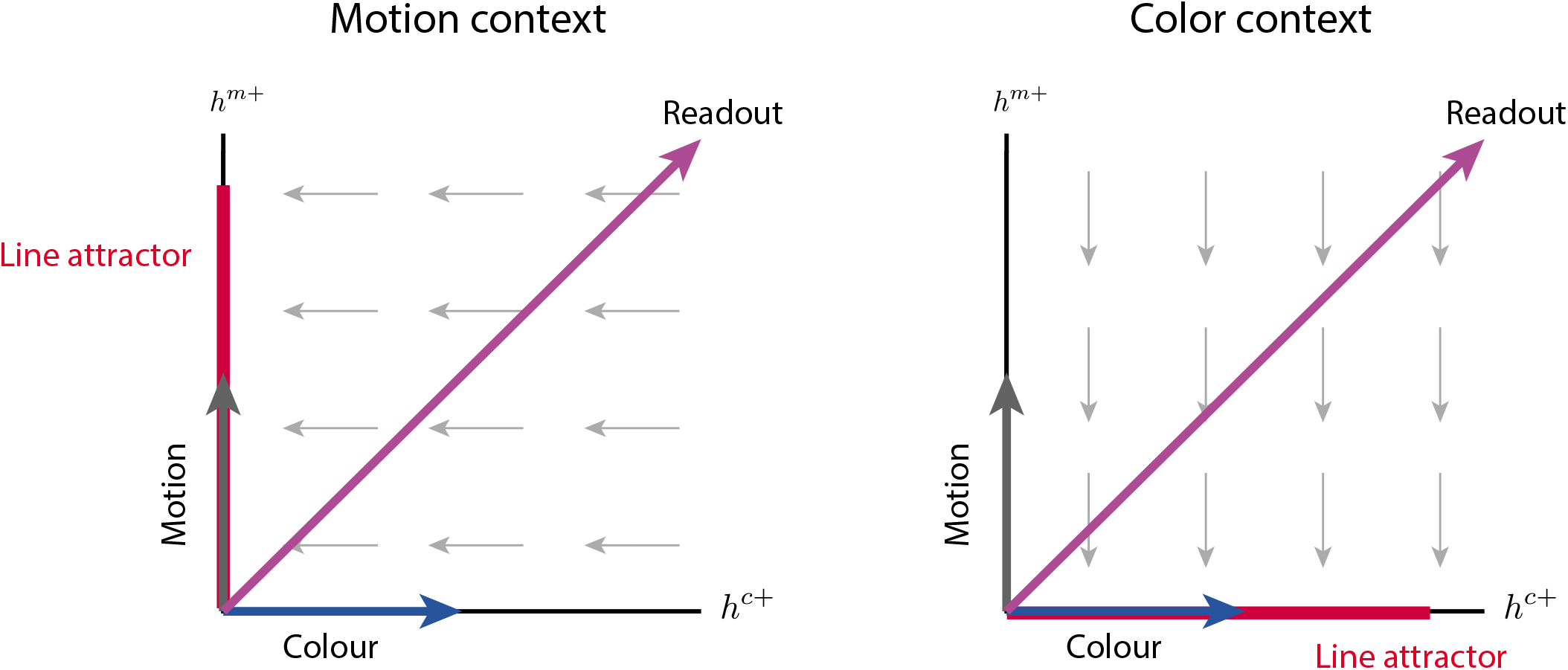
Phase portrait. In both contexts (left and right panels), motion and colour inputs as well as output readout lie in identical directions. However, the dynamics flow against the colour input in the motion context, and against the motion input in the colour context, eventually selecting the appropriate stimulus dimension (‘late selection’). In each context, a line attractor sits on the relevant stimulus dimension.

